# Location of NMDARs and co-agonist control the generation of bursts in nigral dopamine neurons

**DOI:** 10.1101/2024.06.06.597701

**Authors:** Sofian Ringlet, Zoraide Motta, Laura Vandries, Vincent Seutin, Kevin Jehasse, Laura Caldinelli, Loredano Pollegioni, Dominique Engel

## Abstract

NMDA receptor activation in pars compacta substantia nigra dopamine neurons is central to the generation of bursting activity, a key signal temporally associated to movement initiation. The site of the NMDAR pool (synaptic and/or extrasynaptic) as well as the identity of the co-agonist involved in the ignition of this phasic activity remains unknown. Using *ex vivo* electrophysiological recordings, we demonstrate that NMDARs located outside synapses are preponderant for this firing. This pool of receptors is recruited during intense synaptic activity via spillover of glutamate and require the binding of NMDAR co-agonist glycine for their full activation. Synaptic NMDARs are not directly involved in bursting and are activated by D-serine, a distinct co-agonist. Location dependency of NMDARs and co-agonist underlying burst generation may serve as a guideline in understanding the physiological role of dopamine neurons in health and disease.

**Teaser:** Extrasynaptic NMDARs recruited by spillover and glycine allow the generation of bursts in dopamine neurons.

## INTRODUCTION

Activity of dopamine (DA) neurons in the substantia nigra pars compacta (SNc) provides essential signal to the dorsal striatum for the control of movement (*1*, *2*). The fragility of DA neurons located specifically in this brain region is at the origin of the severe motor deficits expressed in Parkinson’s disease (PD) (*3*). The fundamental function ensured by SNc DA neurons is encoded by transitions of their output signal from spontaneous low-frequency action potential (AP) firing to high-frequency bursts of APs. Bursting is a key electrophysiological signal producing an increase of DA release in postsynaptic target areas (*4*, *5*), which is temporally correlated to locomotion initiation (*1*, *6*). Induction of burst firing is operated by N-methyl-D-aspartate receptors (NMDARs) activated by glutamatergic afferents (*7*) originating from several brain areas but predominantly from the subthalamic nucleus (STN), pedunculopontine nucleus (PPN) and cortex (*8*). A substantial fraction of NMDARs in DA neurons are composed of GluN2B and GluN2D subunits (*9*) as in subthalamic (*10*) and hippocampal interneurons (*11*) generating slow deactivating excitatory postsynaptic currents (EPSCs). These subunits are also commonly expressed at extrasynaptic sites (*12–15*) and form triheteromeric GluN1/GluN2B/GluN2D receptors as in SNc DA neurons at both synaptic and extrasynaptic sites (*16*). While bursts critically depend on the presence of the GluN1 subunit (*17*, *18*), the link between the activation of NMDARs and the generation of a burst is not well understood. This lack of information is surprising since bursts mediate the increased dopamine release which is correlated to movement initiation, a sequence of events that is malfunctioning in the context of PD. Bursting activity is observed in vivo (*19*) but also in *ex vivo* tissue by exogenous applications of NMDA in combination to the SK channel blocker apamin (*20–22*), by glutamate applications or synaptic stimulation (*23*).

In addition to glutamate, a co-agonist such as glycine (*24*) or D-serine (*25*) is an absolute requirement for NMDAR channel opening (*26*). While both amino acids have been shown to serve as endogenous agonists at the NMDAR glycine site, the contribution of the one or the other is dependent on neuronal development (*27*), expression of specific GluN2 subunits (*27*, *28*) and level of synaptic activity (*29*). D-serine is generally associated to the synaptic site where GluN2A-containing NMDARs are preferentially expressed, and glycine to the extrasynaptic site where GluN2B-containing NMDARs are present (*28*). The source of D-serine and glycine could potentially be neuronal (*30*, *31*) and astroglial, respectively, but this is still under debate. The amount of co-agonists in and around the synaptic cleft is tightly controlled by their corresponding synthesis and degrading enzymes and by their specific transporters (*32*). In SNc DA neurons, both EPCSs and bursts can be potentiated by exogenous co-agonists or by the blockade of glycine uptake (*21*) indicating that the co-agonist site of NMDARs is not saturated. In addition, blockade of the co-agonist binding-site on NMDARs abolished bursts, revealing the importance of the co-agonist in this firing pattern (*21*). However, the identity of the endogenous co-agonist as well as the population of NMDARs involved in burst generation remain unknown.

In this study, we determined the location of the NMDAR pool and the identity of the co-agonist implicated in the generation of bursting activity in SNc DA neurons. We found that distinct co-agonists activate synaptic and extrasynaptic NMDARs, although both of these NMDARs populations are composed of the same set of GluN2 subunits. Interestingly, the pool of NMDARs implicated in bursting and the pool of extrasynaptic NMDARs require the same co-agonist, indicating that both pools of NMDARs are identical. These findings shed light on the importance of the location of both the NMDAR pool and the co-agonist implicated in the generation of bursts in SNc DA neurons.

## RESULTS

### The co-agonist glycine activates NMDARs to regulate bursting activity

Bursting activity in SNc DA neurons relies on the tonic activation of NMDARs (*33*) through the binding of the neurotransmitter glutamate. In addition to this main agonist, a co-agonist such as glycine or D-serine participates to the activation of NMDARs as the blockade of the co-agonist site abolishes bursts (*21*). However, which endogenous co-agonist is predominant to activate NMDARs during this phasic activity remains unknown. To identify the co-agonist activating NMDARs implicated in burst generation, enzymatic depletion of the co-agonists was combined to patch-clamp recordings in acute brain slices. DA neurons were identified in current- and voltage-clamp and using post hoc immunohistochemistry for tyrosine hydroxylase (**fig. S1**). Bursting activity was induced by bath application of NMDA and apamin and hyperpolarizing current injection (*21*). In these conditions, 100% of neurons (22 out of 22 neurons) exhibited burst firing. Slices were incubated for 45 minutes with either D-amino acid oxidase (DAAO) (**Fig. 1A**) or glycine oxidase (GO) (**Fig. 1B**) to degrade D-serine or glycine, respectively. While the application of DAAO did not affect the profile of the interspike interval (ISI) distribution, burst duration, number of APs in a burst and burst frequency (**Fig. 1, C to F; Table S1**), the incubation with GO strongly modified the profile of the ISI distribution (**Fig. 1G)** and reduced burst frequency but without changing burst duration and the number of AP/burst (**Fig. 1, H to J; Table S1**). Adding D-serine (100 µM) to DAAO-treated samples modified slightly the ISI distribution profile, increased the depolarizing envelope duration and the number of APs riding on a burst without changing the burst frequency (**Fig. 1, C to F; Table S1).** Supplementing glycine (1 mM) to GO-treated samples modified the ISI distribution profile, increased the depolarizing envelope duration and the number of APs riding on a burst but did however not restore the frequency observed in control (**Fig. 1, G to J; Table S1**). In the absence of the enzymes, the bursting activity remained stable over 45 minutes (**fig. S2; Table S2**). These results indicate that glycine regulate the occurrence of bursts via the activation of NMDARs.

**Fig. 1.**
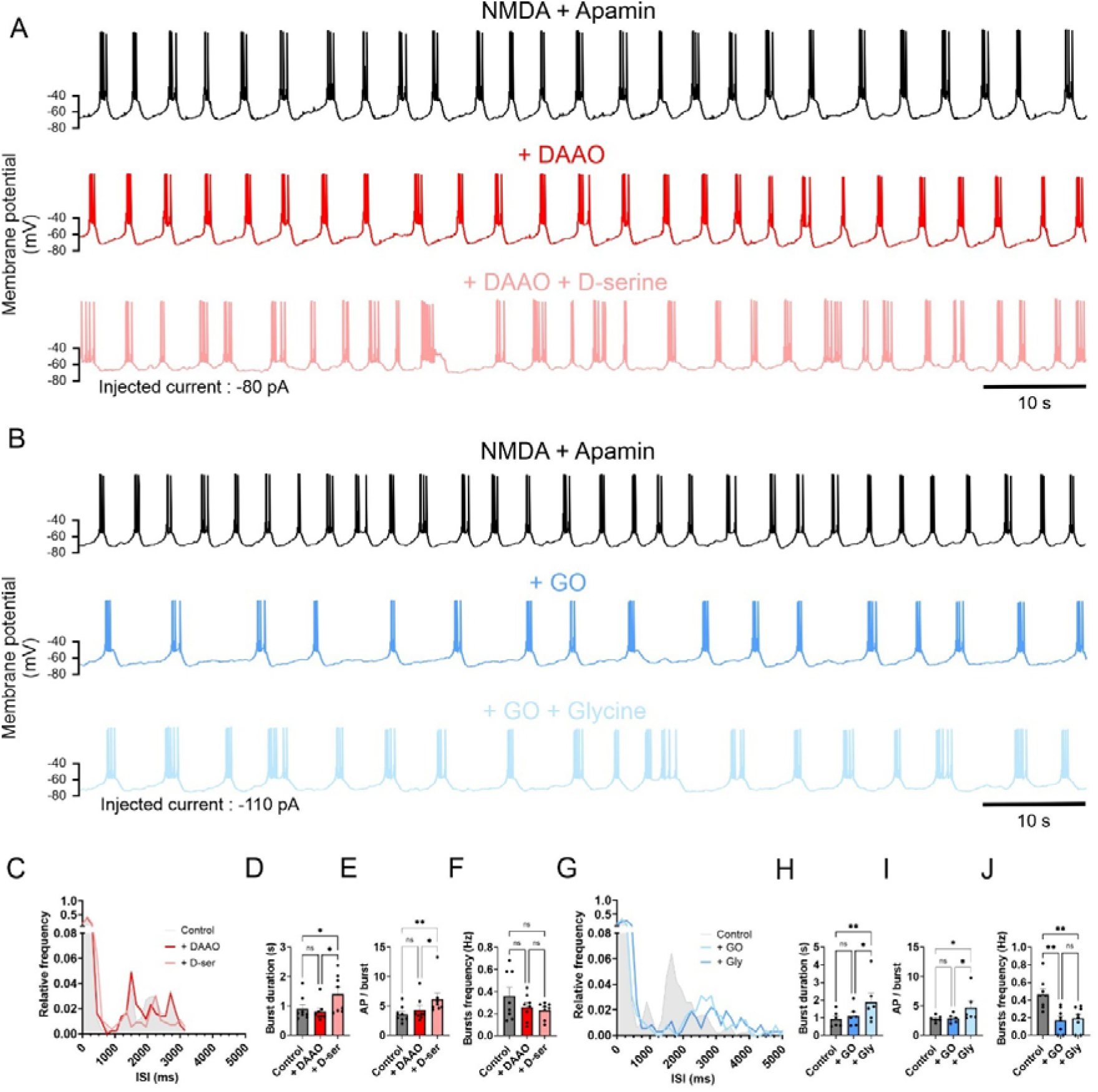
Glycine controls the occurrence of spontaneous bursting. (**A**), Voltage traces from a SNc DA neuron in the presence of NMDA (30 μM) and apamin (300 nM, top trace black), following incubation with DAAO (45 min, 0.2 units/ml; middle trace red) and with DAAO supplemented by D-serine (100 µM, bottom trace light red). Individual neurons were submitted sequentially to DAAO and then to DAAO+D-serine. (**B**), Voltage traces in the presence of NMDA (30 μM) and apamin (300 nM, top trace black), following treatment with GO (45 min, 0.2 units/ml; middle trace blue) and with GO supplemented by glycine (1 mM, bottom trace light blue). Individual neurons were submitted sequentially to GO and then to GO+glycine. (**C**), Histogram for the distribution of interspike interval (ISI) for a DA neuron in control (light grey), in DAAO (red) and in DAAO+D-serine (light red). Note the similar distributions for the three conditions. (**D**), Summary plot showing mean burst duration. Control: 0.90 ± 0.15 s; DAAO: 0.81 ± 0.12 s; DAAO+D-serine: 1.41 ± 0.23 s. Duration was not changed in DAAO (Friedman test followed by post hoc Dunn’s test, ns versus control). Burst duration was increased in DAAO+D-serine (Friedman test followed by post hoc Dunn’s test, \**P* < 0.05 for DAAO+D-serine versus DAAO and for DAAO+D-serine versus control). (**E**), Summary plot showing mean number of action potentials per burst. Control: 3.61 ± 0.48; DAAO: 4.34 ± 0.72; DAAO+D-serine: 6.16 ± 1.05. AP number was not changed in DAAO (Friedman test followed by post hoc Dunn’s test, ns versus control). AP number was increased in DAAO+D-serine (Friedman test followed by post hoc Dunn’s test, \**P* < 0.05 for DAAO+D-serine versus DAAO and \*\**P* < 0.01 for DAAO+D-serine versus control). (**F**), Summary plot showing mean burst frequency. Control: 0.36 ± 0.08 Hz; DAAO: 0.26 ± 0.04 Hz; DAAO+D-serine: 0.23 ± 0.03 Hz. Burst frequency was changed in neither condition (Friedman test followed by post hoc Dunn’s test, ns versus control, ns for DAAO+D-serine versus DAAO and ns for DAAO+D-serine versus control). (**D-F**), Bars represent mean ± SEM; points indicate data from individual experiments. Data points are in control, n=8; DAAO, n=8; DAAO+D-serine, n=8. (**G**), ISI histogram for the distribution of burst activity in control (grey), in GO (blue) and in GO supplemented by glycine (light blue). Note the shift of the second large peak in GO towards higher intervals in comparison to control. (**H**), Summary plot showing burst duration. Control: 0.92 ± 0.18 s; GO: 1.11 ± 0.23 s; GO+glycine: 1.90 ± 0.51 s. Duration was not changed in GO (Friedman test followed by post hoc Dunn’s test, ns versus control). Burst duration was increased in GO+glycine (Friedman test followed by post hoc Dunn’s test, \**P* < 0.05 for GO+glycine versus GO and \*\**P* < 0.01 for GO+glycine versus control). (**I**), Summary plot showing AP per burst. Control: 2.84 ± 0.22; GO: 2.87 ± 0.29; GO + glycine: 4.71 ± 1.10. AP number was not changed in GO (Friedman test followed by post hoc Dunn’s test, ns versus control). AP number was increased in GO+glycine (Friedman test followed by post hoc Dunn’s test, \**P* < 0.05 for GO+glycine versus GO and for GO+glycine versus control). (**J**), Summary plot showing mean burst frequency. Control: 0.47 ± 0.06 Hz; GO: 0.17 ± 0.05 Hz; GO + glycine: 0.19 ± 0.05 Hz. Burst frequency was decreased in GO (Anova test followed by post hoc Tukey’s test, \*\**P* < 0.01 versus control). Frequency was not further changed in GO+glycine (Anova test followed by post hoc Tukey’s test, ns versus GO). Frequency was changed in GO+glycine in comparison to control (Anova test followed by post hoc Tukey’s test, \*\**P* < 0.01). (**H-J**), Bars represent mean ± SEM; points indicate data from individual experiments. Data points are in control, n=8; GO, n=8; GO+glycine, n=8.

To control for changes in the amount of glycine and D-serine in slices in the presence of either GO or DAAO, concentrations of glycine and D-serine were evaluated using high-performance liquid chromatography (HPLC) experiments. In slices incubated for 45 min with either DAAO (0.2 U mL^-1^) or GO (0.2 U mL^-1^), D-serine and glycine concentration was significantly decreased (35.00 ± 12.97% of control; 42.40 ± 11.01% of control respectively). These results confirm that DAAO and GO specifically degrade D-serine and glycine, respectively (**fig. S3**). The specificity of the enzymes observed here is in line with previous observations in comparable experimental conditions (*27–29*) and indicate that the reduction of burst frequency is due to a selective reduction of the respective co-agonist amount.

### Synaptic NMDARs are activated by the co-agonist D-serine

NMDARs are implicated in synaptic transmission in SNc DA neurons (*34*, *35*) but the nature of the endogenous co-agonist activating the synaptic pool of NMDARs has not been identified for these neurons. In addition, whether the identity of the co-agonist for synaptic NMDARs and for bursting activity is identical is not known. Slices were treated with either DAAO to degrade D-serine (**Fig. 2A**) or with GO to reduce the glycine level (**Fig. 2B**) for 45 min. Treatment with DAAO induced a significant reduction in peak amplitude of the NMDAR-mediated component of sEPSCs at +40 mV. Addition of exogenous D-serine (100 µM) reversed the effects of DAAO on the amplitude of sEPSCs which was similar to the EPSC amplitude in control (**Fig. 2C**). Treatment with GO did not change the amplitude of sEPSCs but supplementation of glycine to GO-treated samples increased the amplitude of sEPSCs which was significantly larger than in control conditions (**Fig. 2D**). Currents at −70 mV, representing principally the activation of AMPARs, were not modified, either in the presence of DAAO or GO (**Fig. 2, C and D**). These results provide evidence that D-serine, but not glycine, maintains synaptic activation of NMDARs in DA neurons under basal synaptic activity. Unexpectedly, the co-agonist activating synaptic NMDARs and NMDARs for bursting differed in their identity. This indicates that the pool of NMDARs implicated in the generation of bursts might be different from the pool of synaptic NMDARs and that those may not be directly implicated in the generation of bursts.

**Fig. 2.**
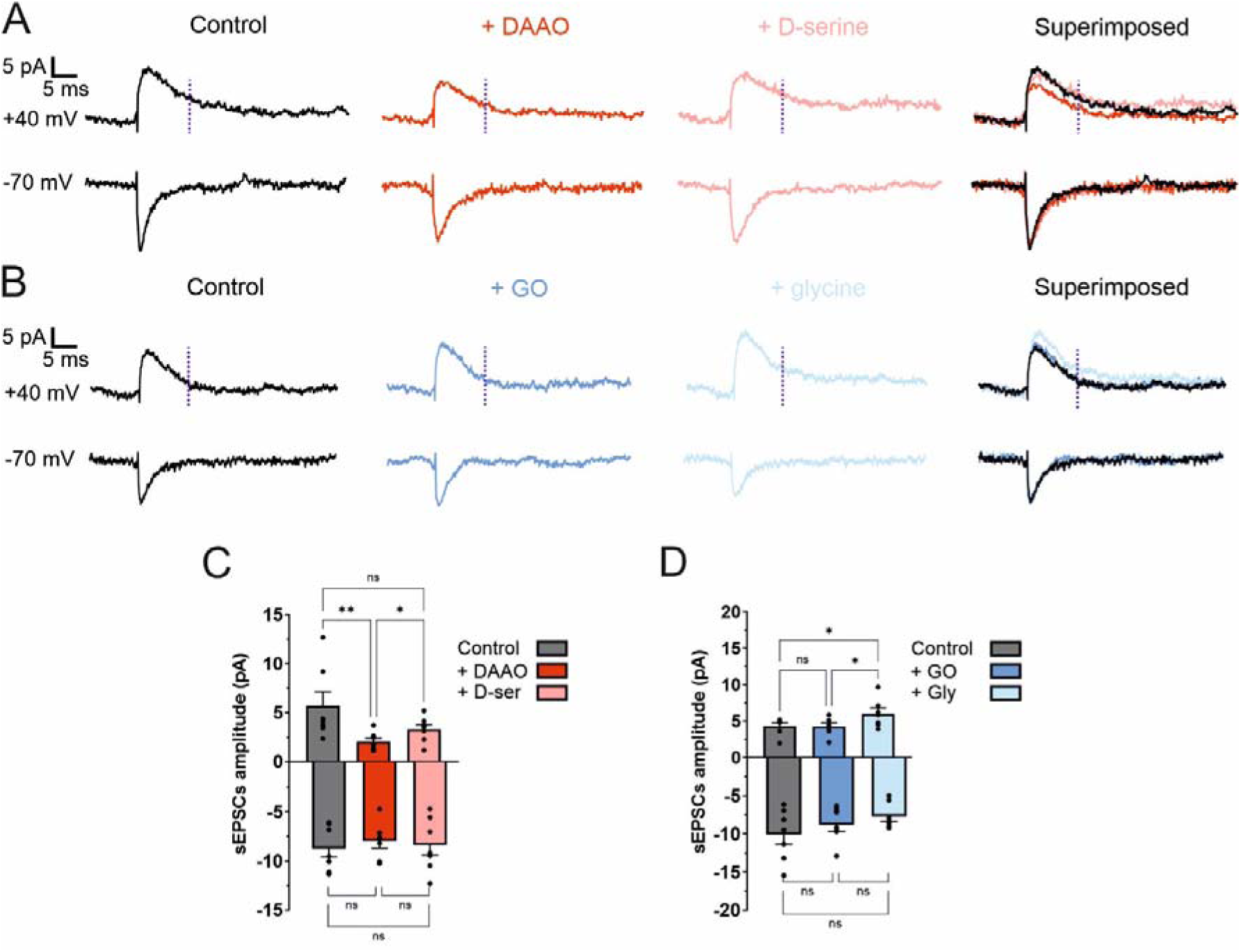
D-serine is the main co-agonist of synaptic NMDA receptors in SNc DA neurons. (**A**), Averaged traces (∼100 events) of sEPSCs at +40 mV (upper traces) and −70 mV (lower traces) in control, in DAAO (45 min, 0.2 units/ml) and in DAAO+D-serine (100 µM). At the right are superimposed traces for the three conditions. For each condition, traces were recorded in the presence of picrotoxin (50 μM), CGP 55845 (1 μM) and strychnine (10 μM) to block GABA_A_, GABA_B_, and Glycine receptors, respectively. Amplitude of NMDAR-mediated component was measured at +40mV (vertical dashed lines). Note the decrease in amplitude of the averaged sEPSC in DAAO at +40 mV. (**B**), Averaged traces (∼100 events) of sEPSCs at +40 mV and −70 mV in control, in GO (45 min, 0.2 units/ml) and in GO+glycine (1 mM). Superimposed traces are at the right. (**C**), Summary bar graph of traces recorded as in (**A**). At +40 mV, control: 5.69 ± 1.43 pA; DAAO: 2.02 ± 0.36 pA; DAAO+D-serine: 3.28 ± 0.49 pA. NMDAR-mediated current was decreased after treatment (Friedman test followed by Dunn’s post hoc test, \*\**P* < 0.01 versus control). NMDAR current was increased in DAAO+D-serine (Friedman test followed by Dunn’s post hoc test, \**P* < 0.05 versus DAAO) but was not changed in comparison to control (Friedman test followed by Dunn’s post hoc test, ns for DAAO+D-serine versus control). At −70 mV, control: −8.84 ± 0.87 pA; DAAO: −8.07 ± 0.71 pA; DAAO+D-serine: −8.48 ± 1.03 pA. AMPAR-mediated currents were changed in neither condition (Anova 1 test followed by Tukey’s post hoc test). Bars represent mean ± SEM; points indicate data from individual experiments. Data points are in control, n=7; DAAO, n=7; DAAO+D-serine, n=7. (**D**), Summary bar graph of traces recorded as in (**B**). At +40 mV, control: 4.30 ± 0.43 pA; GO: 4.26 ± 0.48 pA; GO+glycine: 6.01 ± 0.74 pA. NMDAR current was not changed in GO (Friedman test followed by Dunn’s post hoc test, ns versus control). NMDAR current was increased in GO+glycine (Friedman test followed by Dunn’s post hoc test, \**P* < 0.05 for GO+glycine versus GO and GO+glycine versus control). At −70 mV, control: −10.09 ± 1.29 pA; GO: −8.80 ± 0.86 pA; GO+glycine: −7.69 ± 0.64 pA. AMPAR-mediated currents were changed in neither condition (Anova 1 test followed by Tukey’s post hoc test). Bars represent mean ± SEM; points indicate data from individual experiments. Data points are in control, n=7; GO, n=7; GO+glycine, n=7.

DA neurons in the SNc receive glutamatergic input principally from the STN, the PPN and motor cortex (*8*). Among these inputs, substantial innervation originates from the STN (*8*). To reveal the identity of the co-agonist in this specific synapse, electrically evoked EPSCs (eEPSCs) were recorded in DA neurons during stimulation of STN excitatory afferent fibers. Synaptic transmission originating from the STN was identified using paired-pulse ratio (see Materials and methods and **fig. S4**) and the stimulation intensity was adjusted to slightly above minimal stimulation to activate the smallest number of fibers. Incubation with DAAO reduced significantly the peak amplitude of the NMDAR-mediated component of eEPSCs at +40 mV and addition of exogenous D-serine (100 µM) reversed the effects of DAAO on the amplitude of eEPSCs (**Fig. 3, A and C**). Treatment with GO did not change the mean amplitude of eEPSCs but addition of glycine to GO-treated samples increased the mean amplitude of eEPSCs, which was significantly larger than in control conditions (**Fig. 3, B and D**). Currents at −70 mV were neither changed by the treatment with DAAO, nor with GO. These results obtained for the STN-SNc synapse are similar to those observed on sEPSCs, indicating that the same co-agonist, D-serine, is probably shared by distinct excitatory inputs in DA neurons. In addition, these results further reveal that synaptic NMDARs and NMDAR-mediating bursts are activated by distinct co-agonists.

**Fig. 3.**
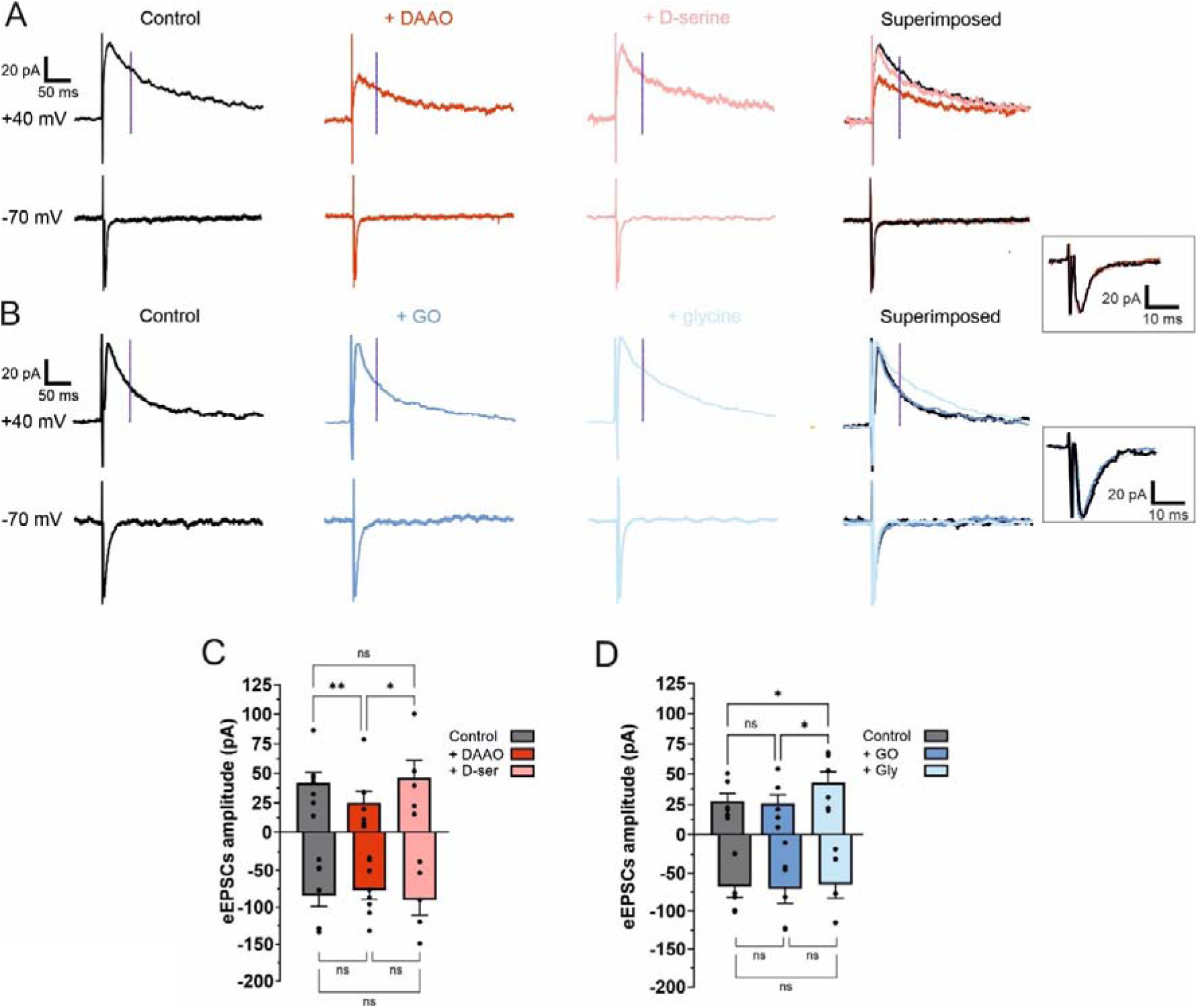
D-serine is the main co-agonist of NMDA receptors in STN-SNc synapses. (**A**), Averaged traces (∼30 events) of eEPSCs at +40 mV (upper traces) and −70 mV (lower traces) in control, in DAAO and in DAAO+D-serine (100 µM). At the right are superimposed traces from the three conditions. A focal double-barreled synaptic stimulating theta electrode was used to activate afferent fibers originating from the STN. For each condition, traces were recorded in the presence of picrotoxin (50 μM), CGP 55845 (1 μM) and strychnine (10 μM). Amplitude of NMDAR-mediated component was measured at +40mV (vertical dashed lines). In the square at the right is an enlargement of the superimposed currents traces at −70 mV. (**B**), Averaged traces (∼30 events) of eEPSCs at +40 mV (upper traces) and −70 mV (lower traces) in control, in GO and in GO+glycine (1 mM). At the right are superimposed traces from the three conditions. In the square at the right is an enlargement of the superimposed currents traces at −70 mV. (**C**), Summary bar graph of experiments as in (**A**). At +40 mV, control: 42.07 ± 8.70 pA; DAAO, 25.10 ± 9.65 pA; DAAO+D-serine: 45.97 ± 15.06 pA. NMDAR-mediated current was decreased after DAAO treatment (Anova 1 test followed by Tukey’s post hoc test, \*\**P* < 0.01 versus control). NMDAR current was increased in DAAO+D-serine (Anova 1 test followed by Tukey’s post hoc test, \**P* < 0.05 versus DAAO) but was not changed in comparison to control (Anova 1 test followed by Tukey’s post hoc test, ns for DAAO+D-serine versus control). At −70 mV, control: −85.19 ± 14.46 pA; DAAO: −77.85 ± 12.51 pA; DAAO+D-serine: −90.78 ± 20.55 pA. AMPAR-mediated currents were changed in neither conditions (Anova 1 test followed by Tukey’s post hoc test). Data points are in control, n=7; DAAO, n=7; DAAO+D-serine, n=5. (**D**), Summary bar graph of traces recorded as in (**B**). At +40 mV, control: 27.90 ± 6.28 pA; GO: 25.91 ± 7.21 pA; GO+glycine: 43.17 ± 8.83 pA. NMDAR current was not changed in GO (Anova 1 test followed by Tukey’s post hoc test, ns versus control). NMDAR current was increased in GO+glycine (Anova 1 test followed by Tukey’s post hoc test, \**P* < 0.05 versus GO). NMDAR mediated current in GO+glycine was similar to current in control (Friedman test followed by Dunn’s post hoc test, ns versus control). At −70 mV, control: −67.94 ± 14.34 pA; GO: −71.00 ± 18.98 pA; GO + glycine: −65.05 ± 18.00 pA. AMPAR-mediated currents were changed in neither conditions (Anova 1 test followed by Tukey’s post hoc test). Bars represent mean ± SEM; points indicate data from individual experiments. Data points are in control, n=7; GO, n=7; GO+glycine, n=5.

### Synaptic NMDARs in nigral DA neurons are triheteromers composed of GluN2B and GluN2D subunits

A strong association between the nature of the predominant co-agonist in excitatory synapses and the expression of a specific GluN2 subunit has been previously established (*27*, *28*). D-serine is usually the main co-agonist found to activate synaptic GluN2A-containing NMDARs. In juvenile DA neurons, a high proportion of synaptic NMDARs are GluN1/GluN2B/GluN2D triheteromers and the presence of the GluN2A subunit has not been detected (*9*). Since D-serine is often the co-agonist of GluN2A-containing NMDARs, the subunit composition of NMDARs in DA neurons has been reexamined in adolescent animals in which a potential switch from GluN2B to GluN2A subunits occurring during the second postnatal week (*36*) already happened. Ambient zinc (Zn^2+^) levels exert a tonic inhibition of EPSCs mediated by GluN2A-containing NMDARs due to a high-affinity binding site for Zn^2+^ specifically on this subunit (*37*). To reveal the functional contribution of the GluN2A subunit, we bath-applied the Zn^2+^ chelator tricine (10 mM) to buffer Zn^2+^ (*38*). This treatment did not modify the amplitude of the NMDAR component of eEPSCs at +40 mV (**Fig. 4, A and B**). The subsequent addition of Zn^2+^ (60 µM, corresponding to 300 nM free Zn^2+^) was also unable to modify the amplitude of eEPSCs. The absence of effect of tricine was also observed for applications of longer durations (**fig. S5**). In contrast, application of the antagonist of NMDARs containing GluN2B (Ro25-6981, 1 µM) reduced significantly the amplitude of eEPSCs to ∼60% of control (**Fig. 4, C and D**). Similarly application of the GluN2D-containing NMDAR antagonist DQP-1105 (20 µM) reduced the amplitude of sEPSCs to ∼53% of control (**Fig. 4, E and F**). Weighted decay time constant of EPSCs was not modified by the application of either Ro25-6981 or DQP-1105. The absence of effect on the decay time constant by either Ro25-6981 or DQP-1105 suggests that synaptic NMDARs are essentially composed of a high proportion of GluN1/GluN2B/GluN2D triheteromers making the participation of diheteromeric GluN1/GluN2B and GluN1/GluN2D receptors unlikely (*11*). Taken together, these results indicate that a large fraction of synaptic NMDARs may be GluN1/GluN2B/GluN2D triheteromers in adolescent SNc DA neurons as in juvenile tissue (*9*) and seem to exclude a switch from Glu2B to GluN2A subunits in DA neurons. In excitatory synapses of DA neurons, D-serine activates triheteromeric NMDARs containing GluN2B and GluN2D but not GluN2A.

**Fig. 4.**
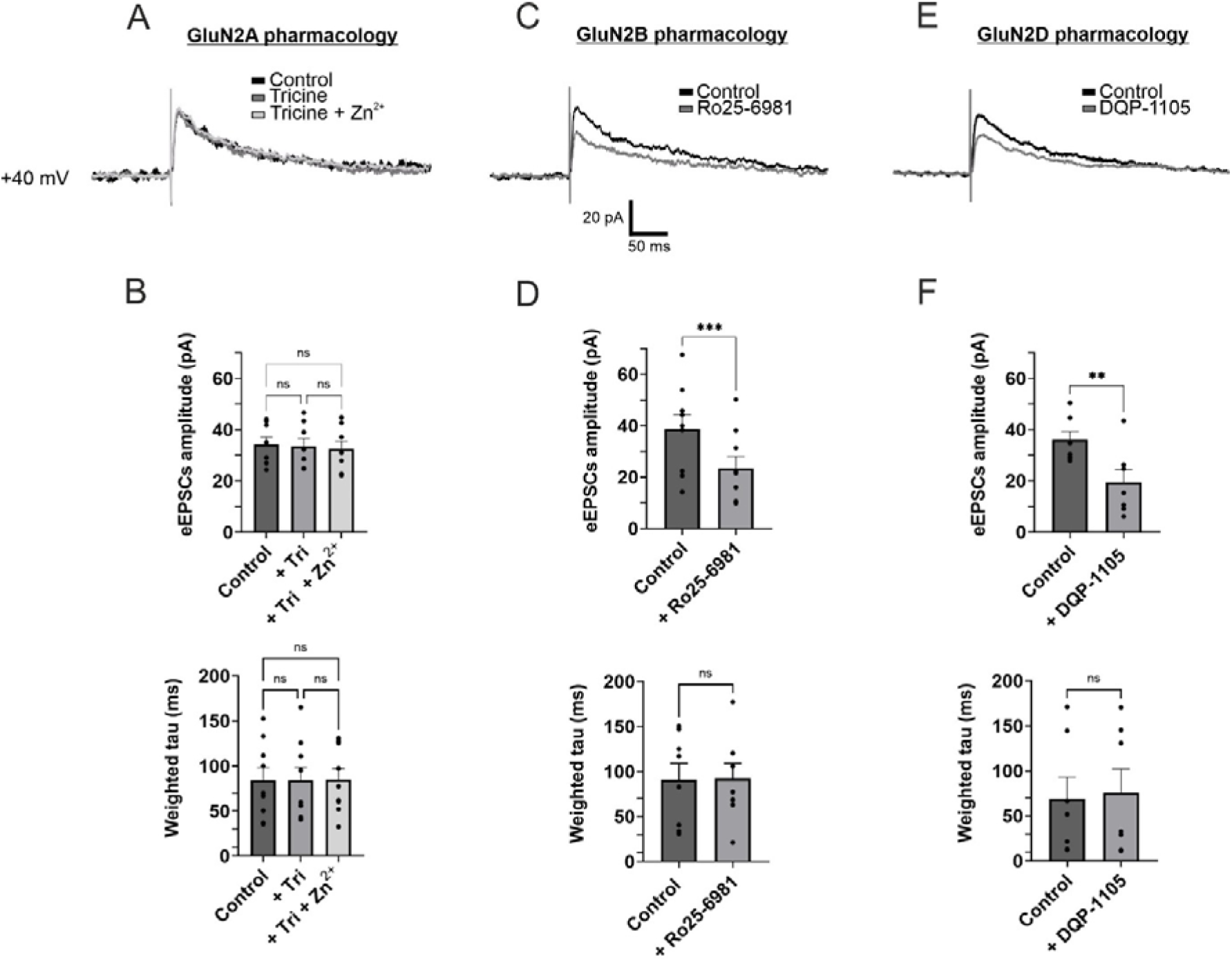
Subunit composition of synaptic NMDARs in nigral DA neurons. (**A**), Averaged eEPSCs (∼30 events) recorded from SNc DA neurons at +40 mV in control conditions (black traces), after incubation with tricine (gray trace) and subsequently after tricine+Zn^2+^ (light grey trace). A focal double-barreled stimulating theta electrode was placed rostrally to the SN to activate fibers from the STN. For each condition traces were recorded in the presence of picrotoxin (50 µM), CGP 55485 (1 µM), CNQX (10 µM), and strychnine (10 µM). tricine was used at 10 mM and Zn^2+^ at 60 µM (to have 300 nM free Zn^2+^). (**B**), Bar chart showing the mean amplitude of NMDAR-eEPSCs (top) in control conditions (dark grey bar), in tricine (grey bar) and in tricine+Zn^2+^ (light grey bar). At +40 mV, control: 34.21 ± 2.80 pA; tricine: 33.48 ± 2.99 pA; tricine+Zn^2+^: 32.40 ± 3.00 pA. NMDAR-mediated current was neither modified in tricine (Anova test followed by Tukey’s post hoc test, ns versus control) nor in tricine+Zn^2+^ (Anova test followed by Tukey’s post hoc test, ns for tricine+Zn^2+^ versus tricine and for tricine+Zn^2+^ versus control). Bar chart showing the weighted decay time constant of the NMDAR-eEPSCs (bottom) in control conditions (dark grey bar), in tricine (grey bar) and in tricine+Zn^2+^ (light grey bar). Weighted decay, control: 84.02 ± 14.13 ms; tricine: 84.01 ± 14.28 ms; tricine+Zn^2+^: 84.99 ± 12.21 ms. Decay of NMDAR current was neither modified in tricine (Anova test followed by Tukey’s post hoc test, ns versus control) nor in tricine+Zn^2+^ (Anova test followed by Tukey’s post hoc test, ns for tricine+Zn^2+^ versus tricine and for tricine+Zn^2+^ versus control). Bars represent mean ± SEM; points indicate data from individual experiments. Data points are in control, n=8; tricine, n=8; tricine+Zn^2+^, n=8. (**C**), Averaged eEPSCs (∼30 events) recorded at +40 mV in control conditions (black traces) and in Ro25-6981 (grey traces). Ro25-6981 was used at 1 µM. (**D**), Bar chart showing the mean amplitude of NMDAR-eEPSCs (top) in control conditions (dark grey bar) and in Ro25-6981 (grey bar). At +40 mV, control: 38.60 ± 5.71 pA; Ro25-6981: 23.42 ± 4.66 pA. NMDAR current amplitude was decreased in Ro25-6981 (Paired T-test, \*\*\**P* < 0.001 versus control). Bar chart showing the weighted decay time constant of the NMDAR-eEPSCs (bottom) in control conditions (dark grey bar) and in Ro25-6981 (grey bar). Weighted decay, control: 91.04 ± 17.93 ms; Ro25-6981: 92.51 ± 16.37 ms). NMDAR current decay was not changed in Ro25-6981 (Paired T-test, ns versus control). Data points are in control, n=9; Ro25-6981, n=9. (**E**), Averaged sEPSCs (∼30 events) recorded at +40 mV in control conditions (black traces) and in DQP-1105 (grey traces,top). DQP-1105 was used at 20 µM. (**F**), Bar chart showing the amplitude of the NMDAR-eEPSCs (top) in control conditions (dark grey bar) and in DQP-1105 (grey bar). At +40 mV, control: 35.99 ± 3.19 pA; DQP-1105: 19.38 ± 4.95 pA). NMDAR current amplitude was decreased in DQP-1105 (Paired T-test, \*\**P* < 0.01 versus control). Bar chart showing the decay of the NMDAR-eEPSCs (bottom) in control conditions (dark grey bar) and in DQP-1105 (grey bar). Weighted decay, control: 68.84 ± 24.38 ms; DQP-1105: 76.05 ± 26.30 ms). NMDAR current decay was not changed in DQP-1105 (Paired T-test, ns versus control). Data points are in control, n=7; DQP-1105, n=7.

The expression of NMDAR subunits in DA neurons was further explored using immunohistochemistry for GluN2A, GluN2B and GluN2D subunits. The distribution of the subunits was determined in TH^+^ DA neurons in the region in which DA neurons were selected for patch-clamp recordings (**fig. S6**). Both GluN2B and GluN2D subunits were present in a large majority (∼80%) of TH^+^ neurons but the GluN2A subunit was only present in ∼20% of TH^+^ neurons (**fig. S6, B and C**). Within TH^+^ neurons, there were ∼3 times more fluorescent dots for GluN2B and GluN2D subunits than for Glun2A subunits (**fig. S6D**). The distribution of the three distinct NMDARs subunits was further examined at higher magnification at the cellular level by taking into account membrane and intracellular locations (**fig. S7**). Immunolabeling for GluN2B and GluN2D subunits was found predominantly at the membrane (GluN2B membrane, ∼75% of the total dots; GluN2D membrane, ∼77% of the total dots) in comparison to the intracellular compartment (GluN2B intracellular, ∼25% of the total dots; GluN2D intracellular, ∼23%). In contrast, the distribution of GluN2A was inverted and found at a high proportion in cell’s intracellular milieu (GluN2A intracellular, ∼85%) and at a very low proportion at the membrane (GluN2A membrane, ∼15%) (**fig. S7, A and B**). This GluN2 subunits expression profile is consistent with the pharmacological data obtained using subunit specific NMDAR blockers (**Fig. 4**).

### Glycine controls the activation of the NMDAR-mediated tonic current

NMDARs are generally concentrated in synapses but are also encountered at extrasynaptic sites (*39*). Activation of extrasynaptic NMDARs by ambient glutamate generates a whole-cell tonic current manifested by the application of the NMDAR blocker D-AP5 (*40*). An NMDAR-mediated tonic current has also been described in SNc DA neurons (*41*) but the identity of the co-agonist activating extrasynaptic NMDARs in SNc DA neurons remains undetermined. At +40 mV, the shift in current produced by D-AP5 had an averaged amplitude of ∼30 pA which is comparable in amplitude to previous reports for the same neurons (*41*) (**Fig. 5, A and B**). In slices incubated with DAAO, the amplitude of the tonic current was not significantly different from control after application of D-AP5. In contrast, in the presence of GO, the amplitude of the tonic current was reduced to ∼46% of control, indicating that reducing the amount of glycine impaired the activation of NMDARs mediating this current. Consistent with previous observations made in other neuronal types (*28*), these results support the hypothesis that glycine is the predominant co-agonist acting at extrasynaptic NMDARs. Interestingly, since glycine is the predominant co-agonist for extrasynaptic NMDARs activated during tonic current and is also controlling the occurrence of bursts, extrasynaptic NMDARs might be critical for the generation of bursts.

**Fig. 5.**
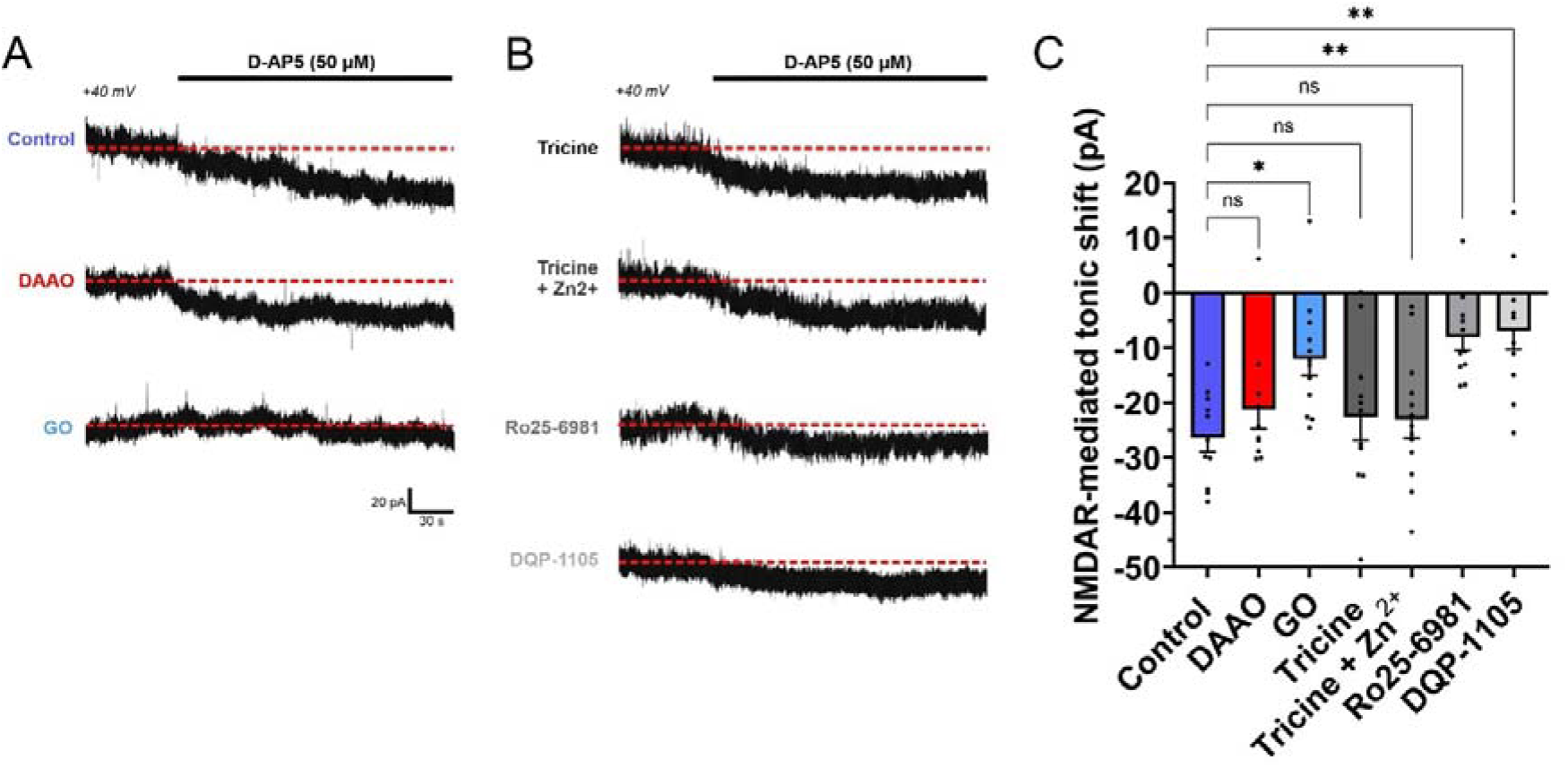
Glycine gates extrasynaptic NMDARs containing GluN2B and GluN2D in nigral DA neurons. (**A**), Tonic NMDAR-mediated current in control conditions (top), during incubation of DAAO (45 min, 0.2 units/ml; below middle) and in GO (45 min, 0.2 units/ml; bottom) revealed by the current shift induced by the application of D-AP5 (50 µM) at a holding potential of +40 mV. Dashed red line indicate amplitude of holding current before the application of D-AP5. Tonic currents were recorded in the presence of CNQX (10 µM), picrotoxin (50 µM), CGP 55485 (1 µM) and strychnine (10 µM). Data in the different conditions are from distinct experiments. (**B**), Tonic NMDAR-mediated current in tricine (10 mM, top), in tricine+Zn^2+^ (60 µM, second from top), in Ro25-6981 (1 µM, third from top) and in DQP-1105 (20 µM, bottom) revealed by the current shift induced by the application of D-AP5 (50 µM) at a holding potential of +40 mV. Dashed red line indicate amplitude of holding current without D-AP5. (**C**), Summary bar graph of the mean amplitude of the NMDAR-mediated tonic current. Control: −26.47 ± 2.51 pA; DAAO: −21.26 ± 3.52 pA; GO: −12.06 ± 3.03 pA; tricine: −22.60 ± 4.22 pA; tricine + Zn^2+^: −23.10 ± 3.24 pA; Ro25-6981: −8.09 ± 2.35 pA; DQP-1105: −6.90 ± 3.45 pA. NMDAR-mediated tonic current was unchanged in DAAO, in tricine and in tricine+ Zn^2+^ (Kruskal-Wallis test followed by a post hoc Dunn’s test, ns versus control). NMDAR-mediated tonic current was reduced in GO (Kruskal-Wallis test followed by a post hoc Dunn’s test, \**P* < 0.05 versus control), in Ro25-6981 and in DQP-1105 (Kruskal-Wallis test followed by a post hoc Dunn’s test, \*\**P* < 0.01 versus control). Bars represent mean ± SEM; points indicate data from individual experiments. Data points are in control, n=11; DAAO, n=10; GO, n=12; tricine, n=11; tricine + Zn^2+^, n=13; Ro25-6981, n=11; DQP-1105, n=11.

### NMDAR subunit composition of NMDAR-mediated tonic current

Since glycine is the co-agonist for the activation of extrasynaptic NMDARs, the GluN2B subunit is expected to be a component of NMDARs at this location (*28*). While NMDAR subtypes at synaptic sites generally contain GluN2A subunits, those at extrasynaptic sites contain GluN2B (*42*, *43*) and GluN2D subunits (*12*, *13*). In adolescent SNc DA neurons the subunit composition of extrasynaptic NMDARs is not known. A large proportion could be GluN1/GluN2B/GluN2D triheteromers as suggested for juvenile DA neurons using outside-out patch recordings (*16*). To determine the subunit composition of extrasynaptic NMDARs in DA neurons, subunit specific NMDAR blockers were used. In the presence of tricine, the magnitude of the tonic current was similar as in control conditions (**Fig. 5, B and C**). Adding Zn^2+^ (60 nM) to tricine did not modify the current amplitude too, indicating that the GluN2A subunit may not compose extrasynaptic NMDARs. The GluN2B blocker Ro25-6981 and the GluN2D blocker DQP-1105 reduced significantly the tonic current to ∼31% of control and ∼26% of control, respectively (**Fig. 5, B and C**). These results indicate that extrasynaptic NMDARs may be composed of GluN2B and GluN2D subunits, as observed for juvenile tissue (*16*) and that extrasynaptic, like synaptic sites, may express a high proportion of the same GluN1/GluN2B/GluN2D NMDAR subtype (*9*). In addition, these observations confirm that GluN2B is not enriched in the extrasynaptic membrane and is present both at synaptic and extrasynaptic sites (*43*) at least in SNc DA neurons. Taken together, our results show that despite similar NMDAR subtypes at synaptic and extrasynaptic sites, distinct co-agonists activate NMDARs at synaptic and extrasynaptic sites. The preference of D-serine to act at synaptic NMDARs and glycine at extrasynaptic NMDARs is therefore more probably due to their availability rather than to the expression of a specific GluN2 subunit.

### Activation of extrasynaptic NMDARs generates tonic current

It is generally accepted that the tonic NMDAR-mediated current is largely due to the activation of extrasynaptic NMDARs (*40*). However, tonically activated synaptic NMDARs could also contribute to the tonic current. To evaluate the potential contribution of synaptic NMDARs to the tonic current, we used the irreversible open channel blocker MK-801 to block this pool of receptors (*44–46*). After a 20 min application of MK-801, low frequency stimulation elicited eEPSCs at +40 mV with strongly reduced amplitude to ∼30% of control due to the blockade of synaptic NMDARs (**Fig. 6A**). The consecutive application of D-AP5 (50 µM) induced a tonic current with an amplitude of ∼22 pA that was similar to the amplitude measured without the suppression of synaptic NMDARs (**Fig. 6, A and C**). In contrast, the blockade of both synaptic and extrasynaptic NMDARs using low frequency repetitive afferent fiber stimulation under MK801 incubation at +40 mV reduced strongly the amplitude of the tonic current (**Fig. 6, B and C**). The absence of a change in the tonic current when synaptic NMDARs are blocked indicates that the pool of extrasynaptic NMDARs sustains the tonic NMDAR-mediated current in DA neurons.

**Fig. 6.**
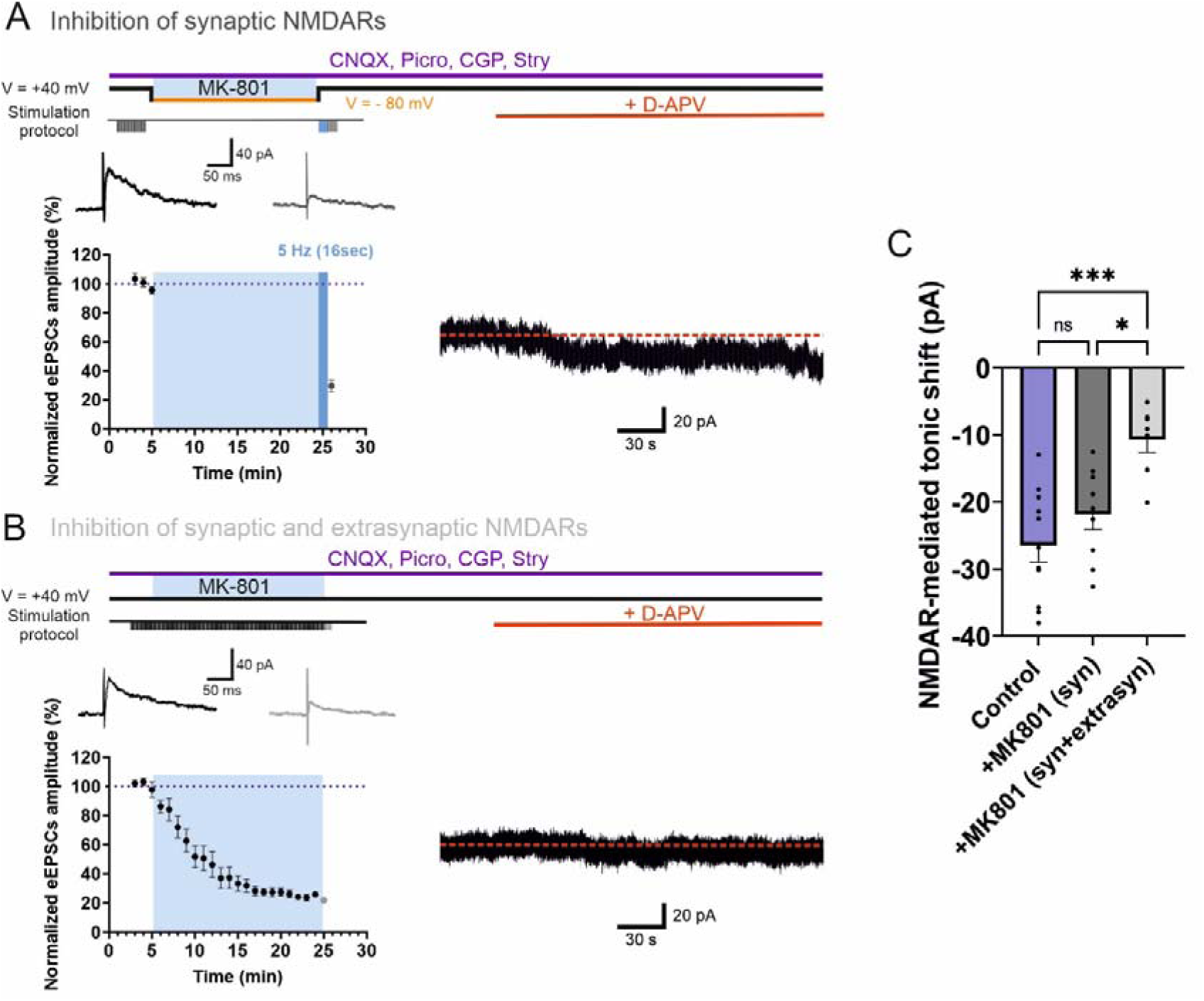
Tonic NMDAR-mediated current is mediated by extrasynaptic NMDARs. (**A**), Experimental protocol to block synaptic NMDARs using the NMDAR open channel blocker MK-801 (left). Control electrically evoked EPSCs (eEPSCs) at V_hold_ = +40 mV under low stimulation frequency (0.1 Hz) in voltage-clamp. Switch to current-clamp (V_hold_ ∼ −80 mV) for ∼20 min during the application of MK-801 (20 µM). Washout of MK-801 and switch back to V_hold_ = +40 mV. Stimulation was resumed using a frequency of 5 Hz for 16 s (*45*, *75*). Note the very strong reduction of the NMDAR-EPSC amplitude when stimulation was resumed. Neuron was kept at V_hold_ = +40 mV and D-AP5 was applied. During the whole experiment, CNQX (10 µM), picrotoxin (50 µM), CGP 55485 (1 µM) and strychnine (10 µM) were present in the bath. (**B**), Experimental design to block both synaptic NMDARs and extrasynaptic NMDARs using MK-801 (left). During application of MK-801, holding potential was kept at +40 mV and repetitive afferent fiber stimulation frequency was low (0.2 Hz). Note the quasi absence of the tonic current after application of D-AP5. (**C**), Summary bar graph of the peak amplitude of the tonic NMDAR-mediated current. Control (without MK-801 treatment): −26.47 ± 2.51 pA; after MK-801 treatment to reduce synaptic NMDARs (syn): −21.84 ± 2.30 pA; after MK-801 treatment and continuous synaptic stimulation to block synaptic and extrasynaptic NMDARs (syn+extrasyn): −10.64 ± 1.97 pA). Tonic NMDAR-mediated current was not different after MK-801 (Anova test followed by a post hoc Tukey’s test, ns versus control) but was decreased after MK-801 and stimulation application (Anova test followed by a post hoc Tukey’s test, \**P* < 0.05 for MK-801+stimulation versus MK-801 and \**P* < 0.05 for MK-801+stimulation versus control). Bars represent mean ± SEM; points indicate data from individual experiments. Data points are in control, n=11; +MK-801 (syn), n=9; +MK-801 (syn+extrasyn), n=7.

### Bursting activity relies on the contribution of extrasynaptic NMDARs

In vivo, bursting activity in SNc DA neurons results from the tonic activation of NMDARs (*33*). In vitro, spontaneous bursts can be generated by exogenous applications of NMDA and the SK channel blocker apamin but whether synaptic or extrasynaptic NMDARs contribute to this firing activity is not known. To determine the possible participation of synaptic NMDARs to the bursting activity, MK-801 (20 µM) was applied to specifically block this pool of receptors. As shown in the previous recordings, the irreversible open-channel inhibitor reduced strongly the amplitude of eEPSCs at +40 mV to ∼26% of control, indicating that a large fraction of synaptic NMDARs were blocked (**Fig. 7A**). The recording configuration was then switched from voltage-clamp at +40 mV to current-clamp (V_membrane_ ∼ −80 mV) to determine whether the spontaneous bursting activity can be induced. After a short time period (∼10 min) to allow stabilization of the membrane potential, NMDA and apamin were bath-applied in combination with small DC current injections. In these conditions, DA neurons exhibited spontaneous bursting activity, similar to the activity recorded without the blockade of synaptic NMDARs (**Fig. 1, A and 1B and fig. S2A**). These recordings were performed using a K^+^-based pipette internal solution and TEA and 4-AP in aCSF to block a majority of the potassium conductances in order to stabilize the current baseline at +40 mV in voltage-clamp. TEA and 4-AP were only present in the bath for voltage-clamp recordings and were washed out while switching to current-clamp. In these conditions, bursting could be generated in a majority of DA neurons (**Fig. 7A**). Bursts generated in these conditions were similar to those in control conditions with comparable burst duration, number of AP/burst and frequency (**Fig. 7, B to E**). These results strongly suggest that bursts in DA neurons are elicited mainly by the activation of extrasynaptic NMDARs.

**Fig. 7.**
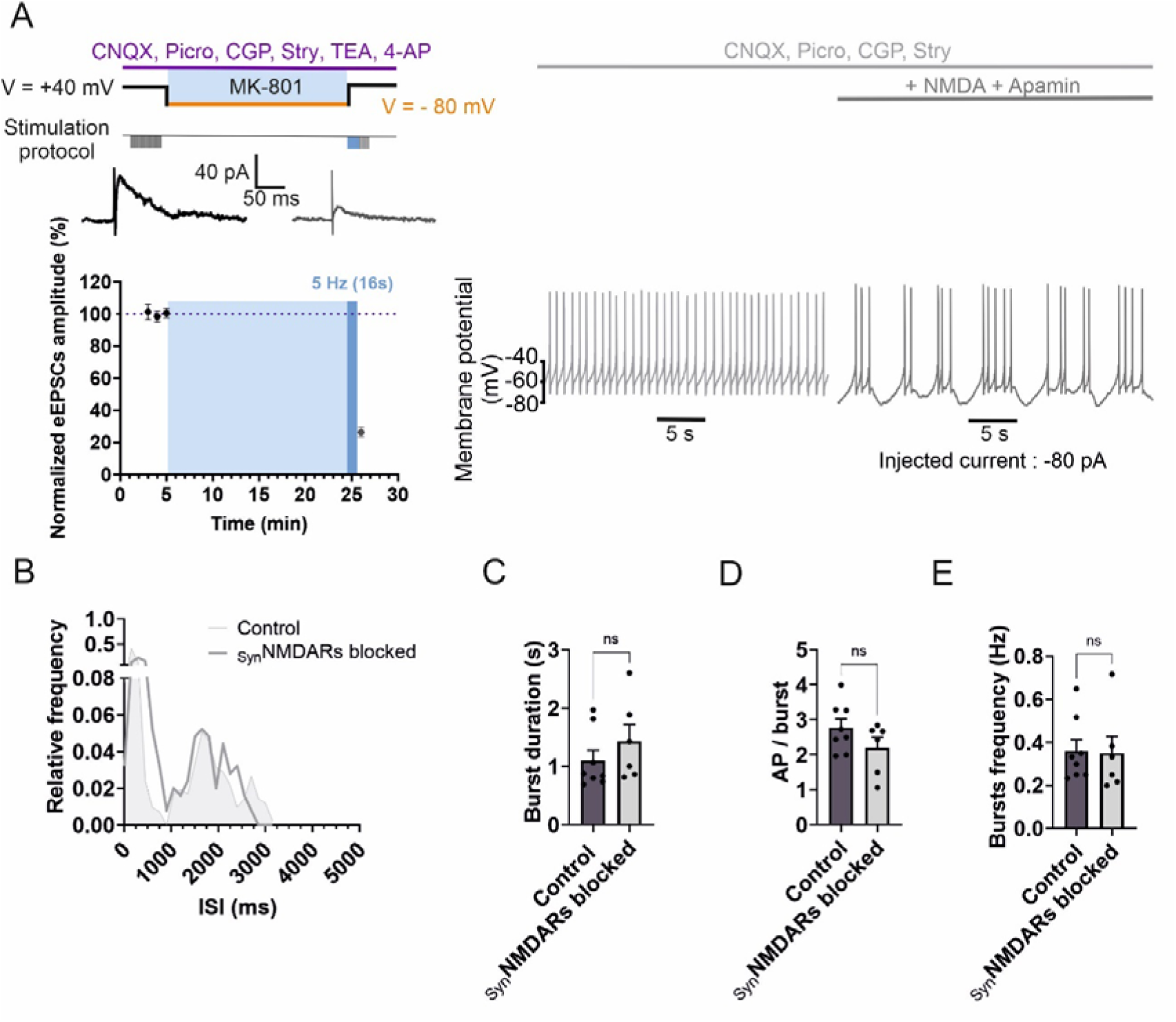
Bursting activity is mediated by extrasynaptic NMDARs. (**A**), Blockade of synaptic NMDARs using MK-801. Control eEPSCs at V_hold_ = +40 mV under low stimulation frequency (0.1 Hz) in voltage-clamp. Switch to current-clamp (V_hold_ = −80 mV) for ∼20 min during the application of MK-801 (20 µM). Washout of MK-801 and switch back to V_hold_ = +40 mV. Stimulation at 5 Hz for 16 s. Bursting activity (right) recorded in current-clamp induced by application of NMDA and apamin. In current-clamp to generate bursts, CNQX (10 µM), picrotoxin (50 µM), CGP 55485 (1 µM) and strychnine (10 µM) were present in the bath. Note the presence of spontaneous bursts after the blockade of synaptic NMDARs. (**B**), Histogram for the distribution of interspike interval (ISI) for a DA neuron in control (light grey) and after blockade of synaptic NMDARs (_syn_NMDARs blocked, dark gray). Note the similar distributions for the two conditions. (**C**), Summary plot showing burst duration in control and after the blockade of synaptic NMDARs. Control: 1.10 ± 0.18 s; _syn_NMDARs blocked: 1.43 ± 0.29 ms. Burst duration was not changed when synaptic NMDARs were blocked (Mann-Whitney test, ns versus control). (**D**), Summary plot showing action potential per burst in control and after the blockade of synaptic NMDARs. Control: 2.77 ± 0.25; _syn_NMDARs blocked: 2.20 ± 0.30. AP number was not changed after synaptic NMDAR blockade (unpaired T-test, ns versus control). (**E**), Summary plot showing burst frequency in control and after the blockade of synaptic NMDARs. Control: 0.36 ± 0.05 Hz; _syn_NMDARs blocked: 0.35 ± 0.08 Hz. Burst frequency was not different after synaptic NMDAR blockade (unpaired T-test, ns versus control). (**C-E**) Bars are mean ± SEM; points indicate data from individual experiments. Data points are in control, n=8; _syn_NMDARs blocked, n=6.

### Synaptic glutamate from STN axons allows burst generation via extrasynaptic NMDARs

Glutamate released during synaptic transmission activates postsynaptic NMDARs in DA neurons to allow the generation of bursts (*23*, *47*). In vivo, the stimulation of the glutamatergic STN neurons favors bursting (*7*). To evaluate the contribution of glutamate released by STN neurons to bursting, fibers originating from this nucleus were stimulated using a focal theta glass electrode and voltage recorded in DA neurons in current-clamp configuration while AMPARs were blocked. Fibers from the STN were identified using paired-pulse facilitation (*48*). In control conditions a train of stimuli at low frequency (10 Hz) for 500 ms did not evoke associated APs (**Fig. 8, A and C to E**). At this stimulation frequency, extrasynaptic NMDARs are not recruited by spillover (*49*). Incubation with either DAAO or GO to reduce D-serine or glycine, respectively, did also not modify the firing. Addition of glycine (1 mM) to GO was not able to change the firing too. In contrast, stimulation at a higher frequency (50 Hz) for 500 ms induced in control conditions and in the presence of DAAO short bursts of APs (**Fig. 8B**) which were similar in both conditions. In the presence of GO, bursts had a reduced number of AP, a decreased intraburst frequency but an increased time to first AP. However, supplementation of glycine to GO induced burst at 50 Hz, increased the AP number, intraburst frequency and decreased time to first AP in comparison to the condition in which GO is alone (**Fig. 8, B and C to E**). At this stimulation frequency (50 Hz), extrasynaptic NMDARs are recruited by spillover of glutamate (*49*) and co-activated by the presence of glycine. These results obtained with synaptically generated bursting corroborate those with spontaneous bursting and further confirm the contribution of extrasynaptic NMDARs and glycine to the generation of bursts.

**Fig. 8.**
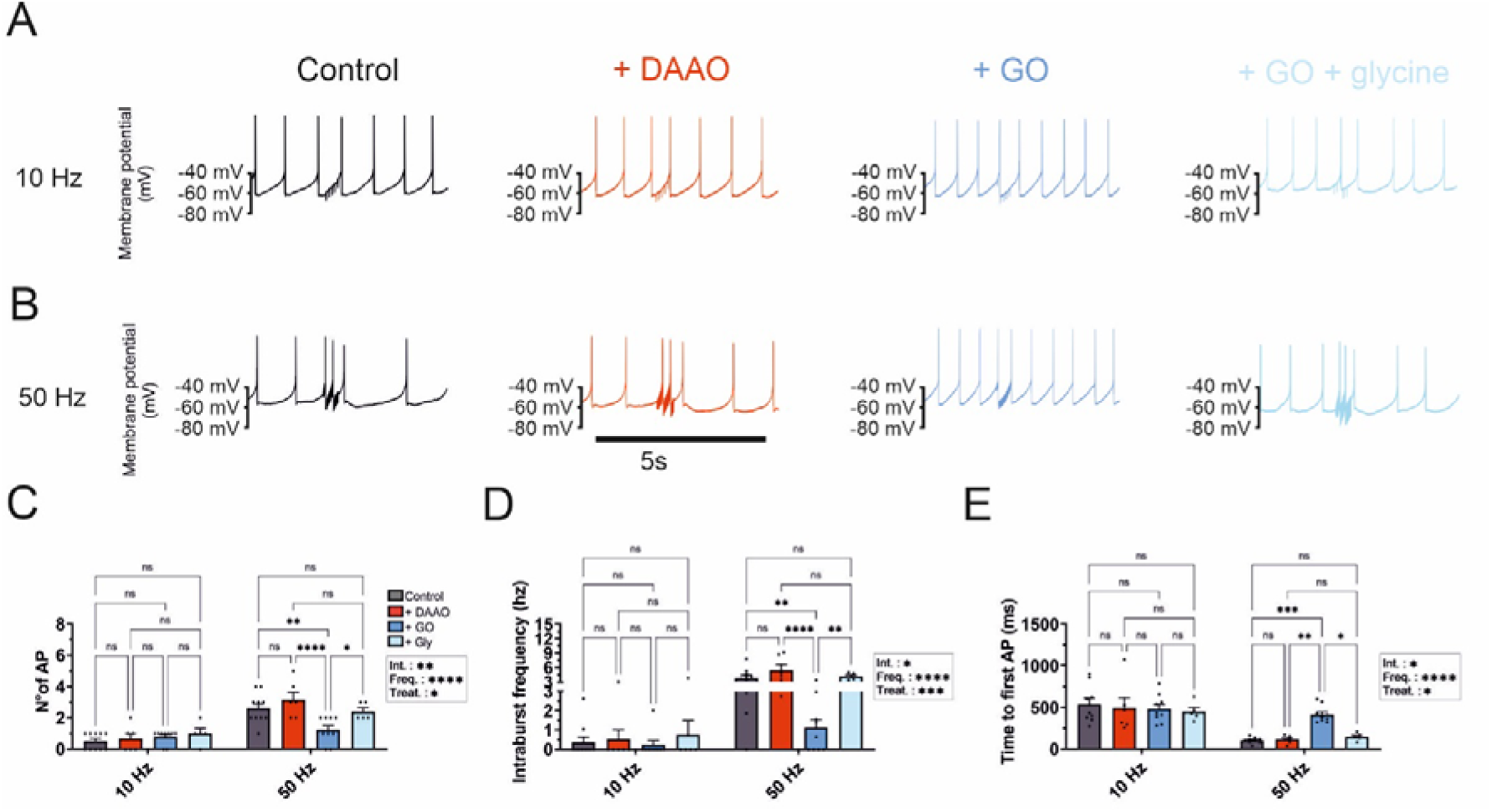
Bursting activity requires glycine and glutamate spillover during high regime of synaptic transmission. (**A**), Voltage traces of action potential in a SNc DA during local synaptic stimulation at 10 Hz for 500 ms in control (left), in DAAO (45min, 0.2 units/ml, middle left), in GO (45min, 0.2 units/ml, middle right) and in GO (45min, 0.2 units/ml) + glycine (1 mM, right). Voltage traces were recorded in the presence of CNQX (10 µM), picrotoxin (50 µM), CGP 55485 (1 µM) and strychnine (10 µM). (**B**), Voltage traces of action potential in a SNc DA during local synaptic stimulation at 50 Hz for 500 ms in control (left), in DAAO (middle left), in GO (middle right) and in GO + glycine (1 mM, right). (**C**), Summary plot showing the mean number of action potential generated during local synaptic stimulation at 10 Hz (left). Control: 0.50 ± 0.17; DAAO: 0.67 ± 0.33; GO: 0.78 ± 0.15; GO + glycine: 1.00 ± 0.32. The number of APs was changed in neither condition (Anova 2 followed by Tukey’s post hoc Test, ns). AP number for synaptic stimulation at 50 Hz (right). Control: 2.60 ± 0.31; DAAO: 3.17 ± 0.48; GO: 1.22 ± 0.28; GO + glycine: 2.40 ± 0.25. AP number was not different in DAAO and in GO+glycine (Anova 2 followed by Tukey’s post hoc Test, ns versus control) but was decrease in GO (Anova 2 followed by Tukey’s post hoc Test, \*\**P* < 0.01 for GO versus control and \*\*\*\**P* < 0.0001 for GO versus DAAO). AP number was increased in GO+glycine in comparison to GO (Anova 2 followed by Tukey’s post hoc Test, \**P* < 0.05). (**D**), Summary plot showing the intraburst frequency during local synaptic stimulation at 10 Hz (left). Control: 0.36 ± 0,27 Hz; DAAO: 0.50 ± 0.50 Hz; GO: 0.22 ± 0.18 Hz; GO + glycine: 0.75 ± 0.73 Hz. Intraburst frequency was changed in neither condition (Anova 2 followed by Tukey’s post hoc Test, ns). Intraburst frequency for stimulation at 50 Hz (right). Control: 3.79 ± 0.66 Hz; DAAO, 5.49 ± 1.26 Hz; GO, 1.11 ± 0.44 Hz; GO + glycine, 4.15 ± 0.33 Hz. Intraburst frequency was not different in DAAO and in GO+glycine (Anova 2 followed by Tukey’s post hoc Test, ns versus control) but was decreased in GO (Anova 2 followed by Tukey’s post hoc Test, \*\**P* < 0.01 for GO versus control and \*\*\*\**P* < 0.0001 for GO versus DAAO). Intraburst frequency was increased in GO+glycine in comparison to GO (Anova 2 followed by Tukey’s post hoc Test, \*\**P* < 0.01). (**E**), Summary plot showing the time to the first action potential during local synaptic stimulation at 10 Hz (left). Control: 536.40 ± 104.60 ms; DAAO: 491.62 ± 112.93 ms; GO: 483.43 ± 55.71 ms; GO+glycine: 451.97 ± 49.12 ms. Time to first AP was changed in neither condition (Anova 2 followed by Tukey’s post hoc Test, ns). Time to first AP for stimulation at 50 Hz (right). Control: 104.60 ± 11.77 ms; DAAO: 112.93 ± 19.85 ms; GO: 415.52 ± 33.76 ms; GO+glycine: 146.70 ± 20.61 ms. Time to first AP was not different in DAAO and in GO+glycine (Anova 2 followed by Tukey’s post hoc Test, ns versus control) but was increased in GO (Anova 2 followed by Tukey’s post hoc Test, \*\*\**P* < 0.001 for GO versus control and \*\*\*\**P* < 0.0001 for GO versus DAAO). Time to first AP was decreased in GO+glycine in comparison to GO (Anova 2 followed by Tukey’s post hoc Test, \**P* < 0.05). (**C-E**) Bars are means ± SEM; points indicate data from individual experiments. Data points are in control, n=10; DAAO, n=6; GO, n=9; GO + glycine, n=5.

## DISCUSSION

The contribution of NMDARs located at synaptic or extrasynaptic sites and the availability of co-agonists within and outside synapses strongly influence the excitability of DA neurons and therefore the downstream release of dopamine in the striatum. Here, by using electrophysiological and biochemical experiments, we determined the nature of the co-agonist and the pool of NMDARs implicated in bursts generated by SNc DA neurons. First, by combining enzymatic depletion for distinct co-agonists and whole-cell recordings, we identified the co-agonist controlling the initiation of bursts. Second, we uncovered that the co-agonists acting during glutamatergic synaptic transmission and bursting are distinct. Third, we found that the identity of the co-agonist implicated in the activation of extrasynaptic NMDARs and in bursting are identical. Fourth, we showed that the tonic NMDAR-mediated current is predominantly mediated by extrasynaptic NMDARs. Fifth, bursting activity is supported principally by the contribution of extrasynaptic but not synaptic NMDARs. Sixth, immunohistochemical labeling is consistent with electrophysiological recordings and confirms the presence of GluN2B and GluN2D but not GluN2A subunits at the membrane of DA neurons, indicating that the same NMDAR subtype is expressed both at synaptic and extrasynaptic sites. The same NMDAR subunits GluN2B and GluN2D at both locations might therefore invalidate an influence of a co-agonist on the expression of a specific GluN2 subunit or the latter on the distribution of a co-agonist at a given location. Finally, we concluded that bursting in DA neurons is mediated by the activation of glycine-gated extrasynaptic NMDARs.

### Extrasynaptic NMDAR population mediate bursting in SNc DA neurons

NMDARs located at extrasynaptic versus synaptic sites have been shown to have different subunit compositions and to deserve distinct cellular functions (*50*). In adolescent SNc DA neurons, synaptic NMDARs are composed of GluN2B and GluN2D subunits, an expression pattern already observed at postnatal ages (*9*). NMDAR-EPSCs were reduced in amplitude to ∼60% of control and ∼50% of control by the GluN2B-containing NMDAR blocker Ro25-6981 and the GluN2D-containing NMDAR blocker DQP-1105, respectively. These percentages of inhibition are similar to those reported by previous observations for midbrain neurons (*10*, *51*) and hippocampal interneurons (*11*) where GluN1/GluN2B/GluN2D triheteromers have been suggested to be present. A small proportion of diheteromeric GluN1/GluN2B or GluN1/GluN2D NMDARs might participate to the synaptic current, but the absence of change of the decay time course of NMDAR-eEPSCs in the presence of either Ro25-6981 or DQP-1105 argues against the presence of these diheteromeric NMDARs (*9*, *52*). Diheteromeric GluN1/GluN2B and GluN1/GluN2D NMDARs produce shorter and longer deactivation time constants than triheteromeric receptors, respectively (*11*). Tricine and Zn^2+^ were ineffective to modify NMDAR-EPSC amplitude, indicating the absence of GluN2A in synaptic receptors. Interestingly, some GluN2A subunits were present in a small proportion of SNc DA neurons as revealed by immunohistochemical labelling, but these were mainly located in the intracellular space with only a minor proportion at the cell membrane. At extrasynaptic sites of DA neurons NMDARs are composed of GluN2B and GluN2D subunits as revealed by GluN2 specific NMDAR blockers on the tonic current, probably the same NMDAR subtypes as those at synaptic locations. A substantial proportion of these triheteromeric receptors is also present in juvenile tissue (*16*), ruling out a subunit exchange until adolescence. A substantial proportion of triheteromeric GluN1/GluN2B/GluN2D NMDARs may therefore be present at both synaptic and extrasynaptic sites. The presence of GluN2B and GluN2D subunits is generally observed at extrasynaptic sites in distinct types of neurons at postnatal or juvenile ages (*12*, *14*, *15*) but also in mature tissue (*13*, *53*). The presence of GluN2B in synaptic and extrasynaptic NMDARs has been also observed in the hippocampus (*43*) suggesting that in this region the subunit present at both sites is mainly in triheteromeric receptors and associated to either GluN2A in synapses (GluN1/GluN2A/GluN2B) NMDARs (*54*) or GluN2D at extrasynaptic locations (GluN1/GluN2B/GluN2D) NMDARs.

### The same NMDAR subtype is activated by distinct co-agonists depending on the location

The identity of the co-agonist has been determined for several glutamatergic synapses in distinct principal neurons and brain regions (*27–29*, *55–57*). D-serine is generally engaged in the activation of synaptic NMDARs and our results confirms this rule for SNc DA neurons. In a unique case, however, glycine has been shown to activate synaptic NMDARs in mature dentate granule cells (*27*). The predominant role of D-serine for synaptic NMDARs has been associated to the presence the GluN2A subunit in postsynaptic NMDARs for instance in adolescent cortical layer 5 neurons (*57*) and hippocampal CA1 pyramidal neurons (*27*, *28*). GluN2A is indeed a component of synaptic NMDARs after the maturational switch from GluN2B to GluN2A (*36*). In the case of DA neurons, our results show that the preponderance of D-serine to activate synaptic NMDARs is not accompanied by the postsynaptic presence of GluN2A but either with the expression of GluN2B and GluN2D in SNc DA neurons. In addition, the developmental switch from GluN2B subunits to GluN2A reported for cortical and hippocampal excitatory synapses (Le Bail et al., 2015) does not seem to occur in SNc DA neurons. Enzymatic reduction of glycine revealed that this co-agonist predominates for the activation of extrasynaptic NMDARs in SNc DA neurons. A comparable correspondence has been reported for CA1 pyramidal neurons where glycine activates also preferentially extrasynaptic GluN2B-containing NMDARs (*28*). The expression of a specific NMDAR subtype (containing GluN2B and GluN2D but not GluN2A) at both the synaptic and extrasynaptic sites challenges the hypothesized association between a given co-agonist and a specific GluN2 subunit expressed in postsynaptic NMDARs (*27*, *28*, *58*). Despite the similar NMDAR subunit composition within and outside synapses, D-serine is active in the synaptic cleft and glycine at extrasynaptic sites, suggesting that a direct influence of the identity of the co-agonist on the presence of a specific GluN2 subtype is unlikely. Since both sites have the same NMDAR subtype composed of GluN2B and GluN2D and D-serine and glycine have similar affinities for a given GluN2 subunit (*58–60*), a discrimination between the two co-agonists cannot be operated by NMDARs. Rather, the organization of the microenvironment, the presence of transporters for the co-agonists and the proximity of astrocytic leaflets in the vicinity of NMDARs certainly influences the predominant role of co-agonists at synaptic and extrasynaptic locations. The glycine transporter GLYT1 expressed in the SNc (*61*) might maintain a high concentration ratio of D-serine/glycine within the synaptic cleft and a low concentration ratio at the extrasynaptic membrane (*62*). D-Serine reuptake by ASCT and EAAT transporters is not so efficient since they are not stereoselective and display low to moderate affinity for D-Serine. The mechanisms for the segregation of glycine and D-serine in the extrasynaptic and synaptic compartments, respectively, are not known despite substantial work dedicated to the distinct functions of the tripartite synapse (*63*). More work is needed to shed light on the establishment and role of the compartmentalization of co-agonists. Interestingly, manipulating the availability of the co-agonists might represent a tool to target specifically synaptic versus extrasynaptic NMDAR pools.

### Contribution of glycine and extrasynaptic NMDARs to bursting activity in SNc DA neurons

Bursting observed in vivo from SNc DA neurons (*19*) can be reproduced in slices by bath application of NMDA and apamin (*20*). The firing in vivo and in vitro is spontaneous and show comparable voltage time courses. The pivotal role of NMDARs in the generation of bursts has been previously revealed in vivo using pharmacological blockade of NMDARs (*33*) and in vitro using genetic inactivation of the essential GluN1 subunit specifically in DA neuron (*17*, *18*). Both of these studies showed that inactivated NMDARs induced a decrease in burst frequency as observed in our work using depletion of glycine. Moreover, glycine is also preponderant for the activation of extrasynaptic NMDARs underlying for a large part the tonic NMDAR-mediated current. These results are in line with work showing that in vivo bursting is generated by tonically activated NMDARs (*33*). Spontaneous bursting activity could be induced after the pharmacological blockade of synaptic NMDARs, further supporting the predominant contribution of extrasynaptic NMDARs to bursting activity. Altogether, these observations strongly suggest that the pool of NMDARs implicated in the generation of bursts and the pool of extrasynaptic NMDARs, both depending on glycine, correspond to the same pool of NMDARs, in line the observations made in vivo (*33*). Functionally, how can extrasynaptic NMDARs be activated to generate bursts? This pool of receptors could be recruited during temporal summation of excitatory synaptic inputs, mediating slow inactivating NMDAR-EPSCs (*18*) due to their slow kinetics (*64*). Extrasynaptic NMDARs are most probably recruited during high frequency synaptic activity by glutamate spillover (*15*) when glutamate uptake is limited or overwhelmed (*41*, *65*). Plateau potentials in layer 5 cortical pyramidal neurons reminiscent of burst in DA neurons are similarly induced by the activation of extrasynaptic NMDARs via spillover (*66*) and hippocampal neurons (*67*, *68*). Extrasynaptic GluN2D-containing NMDARs might favor EPSP-spike coupling, as recently shown for GABAergic neurons (*69*).

Exogenous application of glycine potentiates bursts in SNc DA neurons (*21*) and enzymatic degradation of the co-agonist decreases their frequency as shown here. Glycine is predominant for the activation of extrasynaptic NMDARs which support bursting. These findings represent mechanistic elements that might be considered for pharmacological treatment of PD. Interestingly, stimulating the activation of NMDARs via its co-agonist site could be beneficial during the late phase of PD (*70*). In vivo studies and clinical studies showed indeed that administration of sarcosine, a glycine transporter 1 inhibitor, ameliorates some symptoms of PD (*71*, *72*). Increasing the amount of glycine might act at extrasynaptic NMDARs and favor bursting activity by SNc DA neurons in patients with PD.

## MATERIALS AND METHODS

### Animals

Experiments on Wistar rats were performed in strict accordance with institutional (protocole 2203, Commission d’Éthique Animale, Université de Liège), national (Comité déontologique), and European guidelines for animal experimentation.

### Slice preparation

Adolescent 4- to 7-week-old Wistar rats of either sex were anaesthetized with isoflurane (4% in O_2_, Matrx VIP 3000, Midmark) and subsequently killed by decapitation. Before anesthesia animals were kept in an oxygen-enriched chamber for fifteen minutes in order to increase cell survival. Brain slices were prepared as previously described (*21*). Briefly, after decapitation, the brain was rapidly dissected out and immersed in ice-cold slicing solution containing: 87 mM NaCl, 25 mM NaHCO_3_, 2.5 mM KCl, 1.25 mM NaH_2_PO_4_, 10 mM D-glucose, 75 mM sucrose, 0.5 mM CaCl_2_, and 7 mM MgCl_2_ (pH 7.4 in 95% O_2_/5% CO_2_, ∼325 mOsm). Parasagittal 300 µm-thick (for spontaneous EPSC recordings) or horizontal 250 µm-thick slices (for electrically evoker EPSCS) containing the midbrain were cut using a vibratome (Leica VT-1200, Nussloch, Germany) with a cutting velocity of 0.06 mm/s and a horizontal vibration amplitude of 1.0 mm. Slices containing the SNc were transferred to a storing chamber and allowed to recover at 34°C for 30–45 min. After recovery, slices were kept at room temperature (20 – 25°C) until recording, for up to 4–5 hours.

### Electrophysiology

Slices were transferred to a recording chamber and superfused with aCSF containing (in mM): 125 NaCl, 25 NaHCO_3_, 25 glucose, 2.5 KCl, 1.25 NaH_2_PO_4_, 2 CaCl_2_, and 1 MgCl_2_ (equilibrated with 95% O_2_/5% CO_2_, Osmolarity ∼320 mOsm). DA neurons were visualized using infrared-Dodt gradient contrast (IR-DGC) optics and a 40x objective (numerical aperture 0.75, Zeiss, Oberkochen, Germany) on a Zeiss Axio Examiner.A1 microscope equipped with a CCD camera (C7500-51, Hamamatsu, Japan). The location of the medial terminal (MT) nucleus of the accessory optic tract was used as a reference for the SNc area in horizontal slices. DA neurons were identified in whole-cell using current-clamp and voltage-clamp protocols and post hoc using immunohistochemistry. Selected DA neurons had a voltage rectification with hyperpolarizing current steps, action potentials with long half-width, slow autonomous pacemaking (1-4 Hz) and were positive for tyrosine hydroxylase (**fig. S1**).

Voltage-clamp recordings of DA neurons were conducted with patch pipettes from thick walled borosilicate glass tubing (outer diameter: 2 mm, inner diameter: 1 mm, Hilgenberg, Germany) with a horizontal puller (P-97, Sutter Instruments, Novato, USA). Pipettes were first filled with a potassium gluconate-based internal solution (∼1 mm from the tip) and consecutively with a cesium gluconate solution (*73*) for whole-cell voltage-clamp recordings. For current-clamp recordings, pipettes were only filled with a potassium-based solution. The potassium gluconate solution consisted of, in mM: 120 K-gluconate, 20 KCl, 2 MgCl_2_, 4 Na_2_ATP, 0.5 NaGTP, 5 Na_2_-phosphocreatine, 0.1 EGTA, 10 HEPES and 0.1% biocytin (pH = 7.3; osmolarity: ∼302 mOsm) and the cesium solution: 120 Cs-gluconate, 20 CsCl, 2 MgCl_2_, 4 Na_2_ATP, 0.5 NaGTP, 5 Na_2_-phosphocreatine, 0.1 EGTA, 10 HEPES, 3 QX-314 and 0.1% biocytin (pH = 7.3; osmolarity: ∼306 mOsm). When filled with these solutions, tip resistances were between 3 and 6 MΩ.

Voltage- and current-clamp recordings were obtained using a Multiclamp 700B amplifier (Molecular devices, Palo Alto, CA) connected to a PC via a Digidata 1550 interface (Molecular devices). Bridge balance was used to compensate the series resistance (Rs) in current-clamp recordings (Rs <15 MΩ). The current-clamp signal was sampled at 10 kHz. Whole-cell recordings were discarded if the access resistance increased above 15 MΩ for voltage-clamp and above 20 MΩ for current-clamp or if the values varied over 30% during recording. All voltage-clamp and current-clamp recordings were performed at near-physiological temperature (32-34°C).

Bursting activity in SNc DA neurons was generated by bath-applications of NMDA (30 µM) and apamin (300 nM). Hyperpolarizing current injection (from −80 to −140 pA) was applied to maintain membrane potential in a range (∼-60 to −80 mv) that facilitated the occurrence of bursts. Spontaneous EPSCs were isolated pharmacologically with picrotoxin (50 µM), CGP 55485 (1 µM) and strychnine (10 µM) to block GABA_A_, GABA_B_ and glycine receptor, respectively. Tonic NMDAR-mediated currents were recorded in the presence of CNQX, picrotoxin, CGP 55485 and strychnine. The amplitude of the tonic current was measured by the baseline shift after D-AP5 (50 μM) application.

### Extracellular synaptic stimulation

Afferent fibers were stimulated electrically using theta-glass (borosilicate; Hilgenberg) pipettes (*74*) filled with a HEPES-buffered sodium-rich solution containing (in mM): 135 NaCl, 5.4 KCl, 1.25 NaH_2_PO_4_, 1.8 CaCl_2_, and 1 MgCl_2_, 5 HEPES (pH adjusted to 7.2 with NaOH). The stimulation pipette was placed rostrally close (within ∼100 µm) to the recorded DA neuron and short current pulses were applied (200-600 µA, 200-400 ms) to be slightly above minimal stimulation. Excitatory afferent fibers originating from the STN were selected using paired-pulse ratio. Paired-pulse facilitation was used to identify fibers from the STN as reported previously (*48*). Electrically evoked EPSCs were isolated pharmacologically with GABA_A_, GABA_B_ and glycine receptor blockers (picrotoxin, CGP 55485 and strychnine, respectively).

Synaptic NMDARs were selectively blocked by using the irreversible NMDAR antagonist MK-801 (20 µM) applied for 20 min. Electrically evoked EPSCs (eEPSCs) were recorded at a holding potential of V_hold_ = +40 mV under 0.1 Hz stimulation until stable baseline was obtained. Recording configuration was switched to current-clamp with V_membrane_ ∼ −80 mV for 20 min during the bath-application of MK-801 and stimulation was stopped. MK-801 was washed out and recording configuration was switched back to voltage-clamp with V_hold_ = +40 mV. Stimulation was resumed and delivered at 5 Hz for 16 s to block selectively synaptic NMDARs (*45*, *46*, *75*). Stimulation <10 Hz does not induce measurable extrasynaptic NMDAR-mediated current (*49*).

### Drugs and reagents

Picrotoxin (50 µM, Alomone Lab), CGP 55845 hydrochloride (1 µM, Hellobio), strychnine (10 µM, Sigma) and D-AP5 (50 µM, Alomone labs) were dissolved in doubled distilled water. CNQX disodium (10 µM, Alomone labs) was dissolved in bi-distilled water. Picrotoxin was dissolved in 90% ethanol. Biocytin and ProLong Diamond (Life Technologies). Fluorescein Avidin DCS (Labconsult). The remaining chemicals and all salts were purchased from Sigma-Aldrich (Belgium).

### Immunohistochemistry

Immunohistochemical analysis for identification of DA neurons was done on 300 µm horizontal or parasagittal slices taken after electrophysiological recordings. Biocytin-filled cells were fixed in 4% paraformaldehyde overnight at 4°C. After washing in phosphate buffer saline (PBS, 0.1 M) and then in PBS containing 10% normal goat serum (NGS) and 0.3% triton X-100, slices were incubated with primary antibody against TH (mouse anti-TH 1:1000, Immunostar 22941, Abcam, UK) in PBS containing 5% goat-serum and 0.3% Triton X-100 for 22 hours at room temperature. Subsequently, slices were rinsed with PBS and incubated with the secondary antibody (goat-anti-mouse-Alexa Fluor 568 1:500, Invitrogen A-11004, Thermo Fischer Scientitfic, Waltham, MA, USA) together with Fluorescein Avidin DCS (1 μl/ml, Vector Laboratories A-2001-5,) in PBS and 0.3% triton X-100 overnight at 4°C. After the final rinsing, slices were mounted on a slide containing a 300 µm iSpacer (SunjinLab #IS307, Hsinchu City, Taiwan) with ProLong Gold Antifade (Invitrogen P36934), and imaged using an FV1000 confocal microscope (Olympus, Tokyo, Japan). Images were analyzed with Fiji/ImageJ software (https://imagej.net) and Imaris v9.5.1 (Oxford Instruments, Santa Barbara, CA, USA).

To determine the expression of GluN2 subunits in the SNc immunohistochemical analysis was done on 5- to 6-week-old Wistar rats of either sex. Animals were perfused using a peristaltic pump (Watson Marlow, 101U) with PBS (150 mL), followed by PBS containing with 4% PFA (100 mL). Brains were carefully collected, post-fixed in 4% PFA for 24 hours at 4 °C, subsequently stored in sucrose azide 0.01% for 48 hours at 4°C and sliced (40 µm) with a NX70 cryostat (Epredia Belgium, Machelen, Belgium). Slices were exposed to 10% NGS and 0.1% Triton X-100 followed by primary antibody incubations in PBS containing 5% NGS and 0.1% Triton X-100 overnight at 4 °C. Primary antibodies were against GluN2A subunit (rabbit anti-GluN2A 1:200, Alomone AGC-002) combined to mouse anti-TH (1:1000, Immunostar 22941, Abcam, UK), or GluN2B subunit (rabbit anti-GluN2B 1:200, Alomone AGC-003) with mouse anti-TH or GluN2D subunit (rabbit anti-GluN2D 1:200, Alomone AGC-020) with mouse anti-TH in 5% NGS in PBS and 0.1% Triton X-100 overnight at 4 °C. After rinsing, slices were incubated with secondary antibodies (goat anti-mouse alexa Fluor 568 1:500, Invitrogen A-11004 and goat anti-rabbit Alexa Fluor 488 1:400, Invitrogen A-11011) in PBS with 0.1% Triton X-100 for 2 hours at room temperature. Slices were rinsed with PBS and shortly exposed to Dapi (0.5 µg/mL). After mounting with Aqua-poly/mount (Polysciences 18606, Warrington, PA, USA) images were obtained with an FV1000 confocal microscope (Olympus, Tokyo, Japan) using 5x, 40x and 60x oil immersion objectives (N.A. 0.16, 0.8 and 1.4, respectively).

The distribution of the NMDAR subunits within neurons and at their membranes was analyzed with Imaris using movies (**Movie S1 to S6**). Membrane transparency of reconstructed neurons was adjusted by varying the transparency setting implemented in Imaris to differentiate NMDAR subunits expressed in the cytoplasm (100% transparency - 0% opacity) from NMDAR subunits in the membrane (0% transparency - 100% opacity). TH+ DA neurons were selected around the MT randomly from both hemispheres.

### Enzymes

Recombinant wild-type and M213R variants of *Rhodotorula gracilis* D-amino acid oxidase (RgDAAO; EC 1.4.3.3) and recombinant H244K glycine oxidase from *Bacillus subtilis* (BsGO; EC 1.4.3.19) were produced in *E. coli* cells and purified as previously described (*76–78*). The RgDAAO and BsGO preparations had a specific activity of 75 U/mg protein on D-serine as substrate and 2.5 U/mg protein on glycine as substrate, respectively. Midbrain slices were perfused with RgDAAO (0.2 U/mL) or BsGO (0.2 U/mL) for 45 min. For details about RgDAAO and BsGO substrate specificity see (*77*, *79*).

### High Performance Liquid Chromatography (HPLC) analysis

Slices containing the substantia nigra were stored in the reserve chamber near-physiological temperature (32-34 °C) using three experimental conditions. The first condition was in physiological saline (aCSF), the second in aCSF + GO (0.2 U mL^-1^) and the third in aCSF + DAAO (0.2 U mL^-1^) for 45 minutes. The aCSF collected during all experimental conditions were freeze-dried and then resuspended in 25 mM HCl. Following a neutralization step adding 0.2 M NaOH, the samples were precolumn derivatizated with o-phthaldialdehyde/N-acetyl-L-cysteine in a 1:4 volume ratio of 0.2 M borate, pH 8.2, and 0.2 M sodium phosphate, pH 10. Diastereoisomer derivatives were resolved on a Kinetex EVO C18 reverse-phase column (5 µm, 4.6 x 150 mm; Phenomenex, Bologna, MI, Italy) using a HPLC PU-2089 System (Jasco Europe, Cremella, LC, Italy) equipped with a fluorescence detector (344/443 nm, gain 1000X), in isocratic conditions (0.1 M sodium acetate buffer, pH 6.2, 3% tetrahydrofuran, 0.7 mL/min flow rate), modified from (*80*). Identification and quantification of the D- and L-serine and glycine was based on retention times and peak areas, compared with those associated with external standards (calibration curves were built by injecting 0.005-0.2 pmol of standards). To confirm the identity of D-enantiomers, a selective degradation of D-amino acids in aCSF samples was obtained by adding 10 µg of M213R RgDAAO for 4 h at 30 °C. Statistical analyses were performed using GraphPad Prism 9.0.

### Data acquisition and analysis

Pulse generation and data acquisition were performed using clampEx (clampEx 10.5, Molecular Devices). Data analysis was performed using pClamp 10.5 (clampFit), Mathematica 13 (Wolfram Research, Champaign, IL), Stimfit (https://github.com/neurodroid/stimfit/wiki/Stimfit) and Prism 8 (GraphPad software, La Jolla, CA). To detect sEPSCs, MiniAnalysis (v6.0.7, Synaptosoft Inc.) was used and every single event was inspected and selected visually. To obtain an averaged EPSC, events were superimposed to the initial phase of the current rise. To measure the parameters of the averaged EPSC (amplitude, 20-80% rise time and decay), a short baseline (5 ms) was used immediately preceding the rise of the event. The averaged event was filtered at 2 kHz.

For the analysis of the NMDAR current component of sEPSCs at +40 mV, a dashed vertical line was placed at 10 ms after the onset of the averaged EPSC to evaluate the amplitude of the NMDAR-dependent synaptic current and for eEPSC, it was placed 40 ms after the onset of the event (*29*, *35*). At this time point, the AMPAR-current component is back to baseline. The peak amplitude of the AMPAR-current component was measured at the peak of the averaged EPSC at −70 mV.

Bursts were defined as spontaneously occurring biphasic changes in the membrane potential during which hyperpolarized periods between −60 and −80 mV alternated with depolarized plateau potentials > −60 mV superimposed by APs. This firing activity was stable and reproducible in the presence of NMDA with the SK channel blocker apamin and hyperpolarizing current injection (*20*, *21*).

### Statistical analysis

Experimental values are expressed as mean ± standard error of the mean (SEM). Error bars in the figures also indicate SEM. Statistical analysis was performed using Prism 8. Data were tested for normal distribution with Shapiro-wilk and d’Agostino test. Accordingly, paired-data were either tested by ANOVA 1 test followed by a Tukey post hoc test or Friedman test followed by Dunn’s post hoc when more than 2 conditions. Unpaired-data were tested with Kruskal-Wallis test followed by Dunn’s test. In the case of 2 conditions, either T-test or Wilcoxon test (paired data) and Mann-Withney test (unpaired data) were used. In the case of multiple conditions in 2 different stimulations protocol, ANOVA 2 test followed by a Tukey post hoc test were applied. Significance level is given as * < 0.05, ** < 0.01, *** < 0.001, **** < 0.0001 and ns (not significant) corresponding to > 0.05 is represented in the histograms.

## Acknowledgements

We thank Drs. Jean-Marc Goaillard and for critical reading of the manuscript and helpful comments, Sandy El Sayed and Laurent Massotte for excellent technical assistance and the collaboration with the GIGA-imaging platform for using the confocal microscope and Imaris. **Funding:** This work was supported by grants from the Belgian F.R.S. - F.N.R.S. (FRIA), the Fondation Leon Fredericq, Prix de l’espoir from Marie-José De Ridder and S. R. Fonds facultaires de l’Université de Liège to D.E. Z.M. is a PhD student of the Life Sciences and Biotechnology course at the University of Insubria. L.P. thanks the support from FAR Fondo di Ateneo per la Ricerca.

## Author contributions

Conceptualization: DE

Methodology: DE, LP, SR, KJ, LC, ZM and LV

Investigation: SR, LV, and LC

Writing – Original Draft: DE

Writing – Review & Editing: DE and SR

Funding Acquisition: SR, LP and DE

Resources: VS and LP

Supervision: DE

## Data and materials availability

All data needed to evaluate the conclusions in the paper are present in the paper and/or the Supplementary Materials.

## REFERENCES

1. M. W. Howe, D. A. Dombeck, Rapid signalling in distinct dopaminergic axons during locomotion and reward. Nature 535, 505–510 (2016).

2. J. A. da Silva, F. Tecuapetla, V. Paixão, R. M. Costa, Dopamine neuron activity before action initiation gates and invigorates future movements. Nature 554, 244–248 (2018).

3. O. Hornykiewicz, “The discovery of dopamine deficiency in the parkinsonian brain” in Parkinson’s Disease and Related Disorders (Springer Vienna, Vienna, 2006), pp. 9–15.

4. F. G. Gonon, Nonlinear relationship between impulse flow and dopamine released by rat midbrain dopaminergic neurons as studied by in vivo electrochemistry. Neuroscience 24, 19–28 (1988).

5. R. Ammari, C. Lopez, H. Fiorentino, F. Gonon, C. Hammond, A mouse juvenile or adult slice with preserved functional nigro-striatal dopaminergic neurons. Neuroscience 159, 3–6 (2009).

6. J. E. Markowitz, W. F. Gillis, M. Jay, J. Wood, R. W. Harris, R. Cieszkowski, R. Scott, D. Brann, D. Koveal, T. Kula, C. Weinreb, M. A. M. Osman, S. R. Pinto, N. Uchida, S. W. Linderman, B. L. Sabatini, S. R. Datta, Spontaneous behaviour is structured by reinforcement without explicit reward. Nature 614, 108–117 (2023).

7. K. Chergui, H. Akaoka, P. J. Charléty, C. F. Saunier, M. Buda, G. Chouvet, Subthalamic nucleus modulates burst firing of nigral dopamine neurones via NMDA receptors. [Preprint] (1994).

8. M. Watabe-Uchida, L. Zhu, S. K. Ogawa, A. Vamanrao, N. Uchida, Whole-brain mapping of direct inputs to midbrain dopamine neurons. Neuron 74, 858–873 (2012).

9. S. L. C. Brothwell, J. L. Barber, D. T. Monaghan, D. E. Jane, A. J. Gibb, S. Jones, NR2B- and NR2D-containing synaptic NMDA receptors in developing rat substantia nigra pars compacta dopaminergic neurones. J Physiol 586, 739–50 (2008).

10. S. A. Swanger, K. M. Vance, J. F. Pare, F. Sotty, K. Fog, Y. Smith, S. F. Traynelis, NMDA receptors containing the GluN2D subunit control neuronal function in the subthalamic nucleus. Journal of Neuroscience 35, 15971–15983 (2015).

11. F. Yi, S. Bhattacharya, C. M. Thompson, S. F. Traynelis, K. B. Hansen, Functional and pharmacological properties of triheteromeric GluN1/2B/2D NMDA receptors. Journal of Physiology 597, 5495–5514 (2019).

12. S. G. Brickley, C. Misra, M. H. S. Mok, M. Mishina, S. G. Cull-Candy, NR2B and NR2D subunits coassemble in cerebellar Golgi cells to form a distinct NMDA receptor subtype restricted to extrasynaptic sites. J Neurosci 23, 4958–66 (2003).

13. A. Momiyama, Distinct synaptic and extrasynaptic NMDA receptors identified in dorsal horn neurones of the adult rat spinal cord. Journal of Physiology 523, 621–628 (2000).

14. S. C. Harney, D. E. Jane, R. Anwyl, Extrasynaptic NR2D-containing NMDARs are recruited to the synapse during LTP of NMDAR-EPSCs. Journal of Neuroscience 28, 11685–11694 (2008).

15. N. A. Lozovaya, S. E. Grebenyuk, T. Sh. Tsintsadze, B. Feng, D. T. Monaghan, O. A. Krishtal, Extrasynaptic NR2B and NR2D subunits of NMDA receptors shape ‘superslow’ afterburst EPSC in rat hippocampus. J Physiol 558, 451–463 (2004).

16. S. Jones, A. J. Gibb, Functional NR2B- and NR2D-containing NMDA receptor channels in rat substantia nigra dopaminergic neurones. J Physiol 569, 209–21 (2005).

17. L. P. Wang, F. Li, D. Wang, K. Xie, D. Wang, X. Shen, J. Z. Tsien, NMDA receptors in dopaminergic neurons are crucial for habit learning. Neuron 72, 1055–1066 (2011).

18. L. S. Zweifel, J. G. Parker, C. J. Lobb, A. Rainwater, V. Z. Wall, J. P. Fadok, M. Darvas, M. J. Kim, S. J. Mizumori, C. A. Paladini, P. E. Phillips, R. D. Palmiter, Disruption of NMDAR-dependent burst firing by dopamine neurons provides selective assessment of phasic dopamine-dependent behavior. Proc Natl Acad Sci U S A 106, 7281–7288 (2009).

19. K. Otomo, J. Perkins, A. Kulkarni, S. Stojanovic, J. Roeper, C. A. Paladini, In vivo patch-clamp recordings reveal distinct subthreshold signatures and threshold dynamics of midbrain dopamine neurons. Nat Commun 11, 1–15 (2020).

20. V. Seutin, S. W. Johnson, R. A. North, Apamin increases NMDA-induced burst-firing of rat mesencephalic dopamine neurons. Brain Res 630, 341–344 (1993).

21. G. Destreel, V. Seutin, D. Engel, Subsaturation of the N-methyl-D-aspartate receptor glycine site allows the regulation of bursting activity in juvenile rat nigral dopamine neurons. Eur J Neurosci 50, 3454–3471 (2019).

22. M. E. Soden, G. L. Jones, C. A. Sanford, A. S. Chung, A. D. Güler, C. Chavkin, R. Luján, L. S. Zweifel, Disruption of Dopamine Neuron Activity Pattern Regulation through Selective Expression of a Human KCNN3 Mutation. Neuron 80, 997–1009 (2013).

23. S. N. Blythe, J. F. Atherton, M. D. Bevan, Synaptic Activation of Dendritic AMPA and NMDA Receptors Generates Transient High-Frequency Firing in Substantia Nigra Dopamine Neurons In Vitro. J Neurophysiol 97, 2837–2850 (2007).

24. J. W. Johnson, P. Ascher, Glycine potentiates the NMDA response in cultured mouse brain neurons. [Preprint] (1987). 10.1038/325529a0.

25. N. W. Kleckner, R. Dingledine, Requirement for Glycine in Activation of NMDA-Receptors Expressed in Xenopus Oocytes. Science 241, 835–837 (1988).

26. M. C. Currás, B. S. Pallotta, Single-channel evidence for glycine and NMDA requirement in NMDA receptor activation. Brain Res 740, 27–40 (1996).

27. M. Le Bail, M. Martineau, S. Sacchi, N. Yatsenko, I. Radzishevsky, S. Conrod, K. Ait Ouares, H. Wolosker, L. Pollegioni, J.-M. Billard, J.-P. Mothet, Identity of the NMDA receptor coagonist is synapse specific and developmentally regulated in the hippocampus. Proc Natl Acad Sci U S A 112, E204–13 (2015).

28. T. Papouin, L. Ladépêche, J. Ruel, S. Sacchi, M. Labasque, M. Hanini, L. Groc, L. Pollegioni, J. P. Mothet, S. H. R. Oliet, Synaptic and extrasynaptic NMDA receptors are gated by different endogenous coagonists. Cell 150, 633–646 (2012).

29. Y. Li, S. Sacchi, L. Pollegioni, A. C. Basu, J. T. Coyle, V. Y. Bolshakov, Identity of endogenous NMDAR glycine site agonist in amygdala is determined by synaptic activity level. Nat Commun 4, 1760 (2013).

30. S. Neame, H. Safory, I. Radzishevsky, A. Touitou, F. Marchesani, M. Marchetti, S. Kellner, S. Berlin, V. N. Foltyn, S. Engelender, J. M. Billard, H. Wolosker, The NMDA receptor activation by D-serine and glycine is controlled by an astrocytic Phgdh-dependent serine shuttle. Proc Natl Acad Sci U S A 116, 20736–20742 (2019).

31. J. M. Wong, O. O. Folorunso, E. V. Barragan, C. Berciu, T. L. Harvey, J. T. Coyle, D. T. Balu, J. A. Gray, Postsynaptic Serine Racemase Regulates NMDA Receptor Function. The Journal of Neuroscience 40, 9564–9575 (2020).

32. J.-P. Mothet, M. Le Bail, J.-M. Billard, Time and space profiling of NMDA receptor co-agonist functions. J Neurochem 135, 210–225 (2015).

33. K. Chergui, P. J. Charléty, H. Akaoka, C. F. Saunier, J. -L Brunet, M. Buda, T. H. Svensson, G. Chouvet, Tonic Activation of NMDA Receptors Causes Spontaneous Burst Discharge of Rat Midbrain Dopamine Neurons In Vivo. Eur J Neurosci 5, 137–144 (1993).

34. T. A. Hage, Y. Sun, Z. M. Khaliq, Electrical and Ca2+ signaling in dendritic spines of substantia nigra dopaminergic neurons. Elife 5, 1–26 (2016).

35. M. Jang, K. B. Um, J. Jang, H. J. Kim, H. Cho, S. Chung, M. K. Park, Coexistence of glutamatergic spine synapses and shaft synapses in substantia nigra dopamine neurons. Sci Rep 5, 14773 (2015).

36. C. Bellone, R. A. Nicoll, Rapid Bidirectional Switching of Synaptic NMDA Receptors. Neuron 55, 779–785 (2007).

37. P. Paoletti, P. Ascher, J. Neyton, High-Affinity Zinc Inhibition of NMDA NR1–NR2A Receptors. Journal of Neuroscience 17, 5711–5725 (1997).

38. A. M. Vergnano, N. Rebola, L. P. Savtchenko, P. S. Pinheiro, M. Casado, B. L. Kieffer, D. A. Rusakov, C. Mulle, P. Paoletti, Zinc dynamics and action at excitatory synapses. Neuron 82, 1101–1114 (2014).

39. R. S. Petralia, Y. X. Wang, F. Hua, Z. Yi, A. Zhou, L. Ge, F. A. Stephenson, R. J. Wenthold, Organization of NMDA receptors at extrasynaptic locations. Neuroscience 167, 68–87 (2010).

40. K. Le Meur, M. Galante, M. C. Angulo, E. Audinat, Tonic activation of NMDA receptors by ambient glutamate of non-synaptic origin in the rat hippocampus. J Physiol 580, 373–83 (2007).

41. A. R. Wild, M. Bollands, P. G. Morris, S. Jones, Mechanisms regulating spill-over of synaptic glutamate to extrasynaptic NMDA receptors in mouse substantia nigra dopaminergic neurons. Eur J Neurosci 42, 2633–43 (2015).

42. N. O. Dalby, I. Mody, Activation of NMDA receptors in rat dentate gyrus granule cells by spontaneous and evoked transmitter release. J Neurophysiol 90, 786–97 (2003).

43. A. Z. Harris, D. L. Pettit, Extrasynaptic and synaptic NMDA receptors form stable and uniform pools in rat hippocampal slices. Journal of Physiology 584, 509–519 (2007).

44. J. E. Huettner, B. P. Bean, Block of N-methyl-D-aspartate-activated current by the anticonvulsant MK-801: selective binding to open channels. Proceedings of the National Academy of Sciences 85, 1307–1311 (1988).

45. W. Koh, M. Park, Y. E. Chun, J. Lee, H. S. Shim, M. G. Park, S. Kim, M. Sa, J. Joo, H. Kang, S.-J. Oh, J. Woo, H. Chun, S. E. Lee, J. Hong, J. Feng, Y. Li, H. Ryu, J. Cho, C. J. Lee, Astrocytes Render Memory Flexible by Releasing D-Serine and Regulating NMDA Receptor Tone in the Hippocampus. Biol Psychiatry 91, 740–752 (2022).

46. D. dan Liu, Q. Yang, S. tian Li, Activation of extrasynaptic NMDA receptors induces LTD in rat hippocampal CA1 neurons. [Preprint] (2013). 10.1016/j.brainresbull.2012.12.003.

47. T. A. Hage, Z. M. Khaliq, Tonic Firing Rate Controls Dendritic Ca2+ Signaling and Synaptic Gain in Substantia Nigra Dopamine Neurons. Journal of Neuroscience 35, 5823–5836 (2015).

48. G. Beaudoin, J. Gomez, J. Perkins, J. Bland, A. Petko, C. Paladini, Cocaine selectively reorganizes excitatory inputs to substantia Nigra pars compacta dopamine neurons. Journal of Neuroscience 38, 1151–1159 (2018).

49. A. Z. Harris, D. L. Pettit, Recruiting Extrasynaptic NMDA Receptors Augments Synaptic Signaling. J Neurophysiol 99, 524–533 (2008).

50. G. E. Hardingham, H. Bading, Synaptic versus extrasynaptic NMDA receptor signalling: Implications for neurodegenerative disorders. Nat Rev Neurosci 11, 682–696 (2010).

51. M. T. Harnett, B. E. Bernier, K.-C. Ahn, H. Morikawa, Burst-timing-dependent plasticity of NMDA receptor-mediated transmission in midbrain dopamine neurons. Neuron 62, 826–38 (2009).

52. R. E. Perszyk, J. O. DiRaddo, K. L. Strong, C. M. Low, K. K. Ogden, A. Khatri, G. A. Vargish, K. A. Pelkey, L. Tricoire, D. C. Liotta, Y. Smith, C. J. McBain, S. F. Traynelis, GluN2D-containing N-methyl-D-aspartate receptors mediate synaptic transmission in hippocampal interneurons and regulate interneuron activity. Mol Pharmacol 90, 689–702 (2016).

53. I. Riebe, H. Seth, G. Culley, Z. Dósa, S. Radi, K. Strand, V. Fröjd, E. Hanse, Tonically active NMDA receptors - a signalling mechanism critical for interneuronal excitability in the CA1 stratum radiatum. Eur J Neurosci 43, 169–178 (2016).

54. C. Rauner, G. Köhr, Triheteromeric NR1/NR2A/NR2B receptors constitute the major N-methyl-D-aspartate receptor population in adult hippocampal synapses. Journal of Biological Chemistry 286, 7558–7566 (2011).

55. A. Panatier, D. T. Theodosis, J. P. Mothet, B. Touquet, L. Pollegioni, D. A. Poulain, S. H. R. Oliet, Glia-Derived d-Serine Controls NMDA Receptor Activity and Synaptic Memory. Cell 125, 775–784 (2006).

56. L. Curcio, M. V. Podda, L. Leone, R. Piacentini, A. Mastrodonato, P. Cappelletti, S. Sacchi, L. Pollegioni, C. Grassi, M. D’Ascenzo, Reduced d-serine levels in the nucleus accumbens of cocaine-treated rats hinder the induction of NMDA receptor-dependent synaptic plasticity. Brain 136, 1216–1230 (2013).

57. P. Fossat, F. R. Turpin, S. Sacchi, J. Dulong, T. Shi, J. M. Rivet, J. V. Sweedler, L. Pollegioni, M. J. Millan, S. H. R. Oliet, J. P. Mothet, Glial D-serine gates NMDA receptors at excitatory synapses in prefrontal cortex. Cerebral Cortex 22, 595–606 (2012).

58. J. S. Ferreira, T. Papouin, L. Ladépêche, A. Yao, V. C. Langlais, D. Bouchet, J. Dulong, J. P. Mothet, S. Sacchi, L. Pollegioni, P. Paoletti, S. H. Oliet, L. Groc, Co-agonists differentially tune GluN2B-NMDA receptor trafficking at hippocampal synapses. Elife 6, 1–22 (2017).

59. K. B. Hansen, L. P. Wollmuth, D. Bowie, H. Furukawa, F. S. Menniti, A. I. Sobolevsky, G. T. Swanson, S. A. Swanger, I. H. Greger, T. Nakagawa, C. J. McBain, V. Jayaraman, C. M. Low, M. L. Dell’Acqua, J. S. Diamond, C. R. Camp, R. E. Perszyk, H. Yuan, S. F. Traynelis, Structure, Function, and Pharmacology of Glutamate Receptor Ion Channels. [Preprint] (2021). 10.1124/pharmrev.120.000131.

60. P. E. Chen, M. T. Geballe, E. Katz, K. Erreger, M. R. Livesey, K. K. O’toole, P. Le, C. J. Lee, J. P. Snyder, S. F. Traynelis, D. J. A. Wyllie, Modulation of glycine potency in rat recombinant NMDA receptors containing chimeric NR2A/2D subunits expressed in Xenopus laevis oocytes. Journal of Physiology 586, 227–245 (2008).

61. B. Cubelos, C. Giménez, F. Zafra, Localization of the GLYT1 glycine transporter at glutamatergic synapses in the rat brain. Cereb Cortex 15, 448–59 (2005).

62. J. G. Dopico, T. González-Hernández, I. M. Pérez, I. G. García, A. M. Abril, J. O. Inchausti, M. Rodríguez Díaz, Glycine release in the substantia nigra: Interaction with glutamate and GABA. Neuropharmacology 50, 548–557 (2006).

63. C. Henneberger, L. Bard, C. King, A. Jennings, D. A. Rusakov, NMDA receptor activation: Two targets for two co-agonists. [Preprint] (2013). 10.1007/s11064-013-0987-2.

64. A. Drotos, Y. Herrera, R. Zarb, M. Roberts, GluN2D-containing NMDA receptors enhance temporal integration in VIP neurons in the inferior colliculus. bioRxiv preprint, 1–13 (2023).

65. N. Arnth-Jensen, D. Jabaudon, M. Scanziani, Cooperation between independent hippocampal synapses is controlled by glutamate uptake. Nat Neurosci 5, 325–331 (2002).

66. K. D. Oikonomou, S. M. Short, M. T. Rich, S. D. Antic, Extrasynaptic glutamate receptor activation as cellular bases for dynamic range compression in pyramidal neurons. Front Physiol, 1–22 (2012).

67. M. Herman, C. Jahr, Extracellular glutamate concentration in hippocampal slice. J Neurosci 27, 9736–41 (2007).

68. T. Suzuki, S. Kodama, C. Hoshino, T. Izumi, H. Miyakawa, A plateau potential mediated by the activation of extrasynaptic NMDA receptors in rat hippocampal CA1 pyramidal neurons. Eur J Neurosci 28, 521–534 (2008).

69. L. Yao, Y. Rong, X. Ma, H. Li, D. Deng, Y. Chen, S. Yang, T. Peng, T. Ye, F. Liang, N. Xu, Q. Zhou, Extrasynaptic NMDA Receptors Bidirectionally Modulate Intrinsic Excitability of Inhibitory Neurons. Journal of Neuroscience 42, 3066–3079 (2022).

70. U. Heresco-Levy, S. Shoham, D. C. Javitt, Glycine site agonists of the N-methyl-d-aspartate receptor and Parkinson’s disease: A hypothesis. Movement Disorders 28, 419–424 (2013).

71. C. H. Tsai, H. C. Huang, B. L. Liu, C. I. Li, M. K. Lu, X. Chen, M. C. Tsai, Y. W. Yang, H. Y. Lane, Activation of N-methyl-D-aspartate receptor glycine site temporally ameliorates neuropsychiatric symptoms of Parkinson’s disease with dementia. Psychiatry Clin Neurosci 68, 692–700 (2014).

72. I. Frouni, E. Kim, J. Shaqfah, D. Bédard, C. Kwan, S. Belliveau, P. Huot, [3H]-NFPS binding to the glycine transporter 1 in the hemi-parkinsonian rat brain. Exp Brain Res 242, 1203–1214 (2024).

73. H.-X. Wang, W.-J. Gao, Development of calcium-permeable AMPA receptors and their correlation with NMDA receptors in fast-spiking interneurons of rat prefrontal cortex. J Physiol 588, 2823–38 (2010).

74. A. Kumar, O. Schiff, E. Barkai, B. W. Mel, A. Poleg-Polsky, J. Schiller, NMDA spikes mediate amplification of inputs in the rat piriform cortex. Elife 7 (2018).

75. Q. Yang, G. Zhu, D. Liu, J. G. Ju, Z. H. Liao, Y. X. Xiao, Y. Zhang, N. Chao, J. Wang, W. Li, J. H. Luo, S. T. Li, Extrasynaptic NMDA receptor dependent long-term potentiation of hippocampal CA1 pyramidal neurons. Sci Rep 7, 1–9 (2017).

76. S. Sacchi, S. Lorenzi, G. Molla, M. S. Pilone, C. Rossetti, L. Pollegioni, Engineering the Substrate Specificity ofd-Amino-acid Oxidase. Journal of Biological Chemistry 277, 27510–27516 (2002).

77. V. Job, G. Molla, M. S. Pilone, L. Pollegioni, Overexpression of a recombinant wild-type and His-tagged *Bacillus subtilis* glycine oxidase in *Escherichia coli*. Eur J Biochem 269, 1456–1463 (2002).

78. S. Fantinato, L. Pollegioni, M. Pilone S., Engineering, expression and purification of a His-tagged chimeric D-amino acid oxidase from Rhodotorula gracilis. Enzyme Microb Technol 29, 407–412 (2001).

79. L. Frattini, E. Rosini, L. Pollegioni, M. S. Pilone, Analyzing the d-amino acid content in biological samples by engineered enzymes. Journal of Chromatography B 879, 3235–3239 (2011).

80. D. Punzo, F. Errico, L. Cristino, S. Sacchi, S. Keller, C. Belardo, L. Luongo, T. Nuzzo, R. Imperatore, E. Florio, V. De Novellis, O. Affinito, S. Migliarini, G. Maddaloni, M. J. Sisalli, M. Pasqualetti, L. Pollegioni, S. Maione, L. Chiariotti, A. Usiello, Age-Related Changes in d-Aspartate Oxidase Promoter Methylation Control Extracellular d-Aspartate Levels and Prevent Precocious Cell Death during Brain Aging. The Journal of Neuroscience 36, 3064–3078 (2016).

