## supplementary materials for "Location of NMDARs and co-agonist control the generation of bursts in nigral dopamine neurons"

Sofian Ringlet et al.

Figs. S1 to S7

Tables S1 to S2

Movies S1 to S6

**Other Supplementary Materials for this manuscript include the following:**

Movies S1 to S6

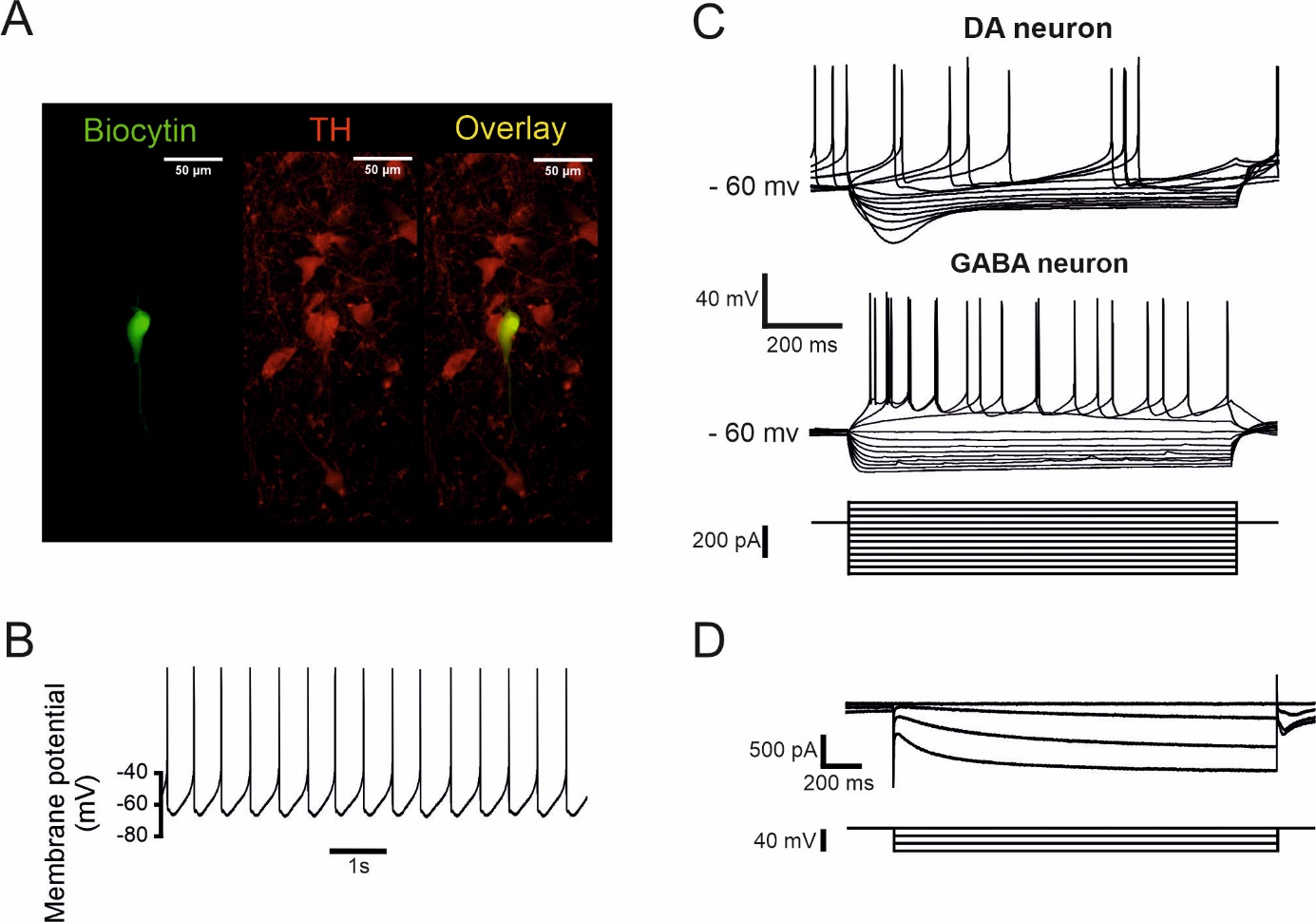

**Fig. S1. Identification of SNc DA neurons**

**(A)** Epifluorescence photomicrographs of a DA neuron filled with biocytin and stained using fluorescein isothiocyanate (FITC)-conjugated avidin (left) and for tyrosine hydroxylase (TH, center). The overlay of FITC and TH is at the right. (**B)** Low frequency (typically ~1 – 4 Hz) spontaneous action potential firing recorded in a DA neuron in current-clamp with no current injection. **(C)** Voltage responses of a whole-cell recorded GABA neuron (top) and a DA neuron (middle) to 1-sec hyper- or depolarizing current pulses (−30 to 70 pA, 10 pA increment, bottom). Note the presence of a rectification in the membrane potential (sag) in the DA neuron in response to a hyperpolarizing current step which is absent in the GABA neuron. **(D)** Current traces in whole-cell voltage clamp in response to hyperpolarizing voltage steps showing slowly increasing currents typical for the activation of hyperpolarization-activated cyclic nucleotide-gated (HCN) channels.

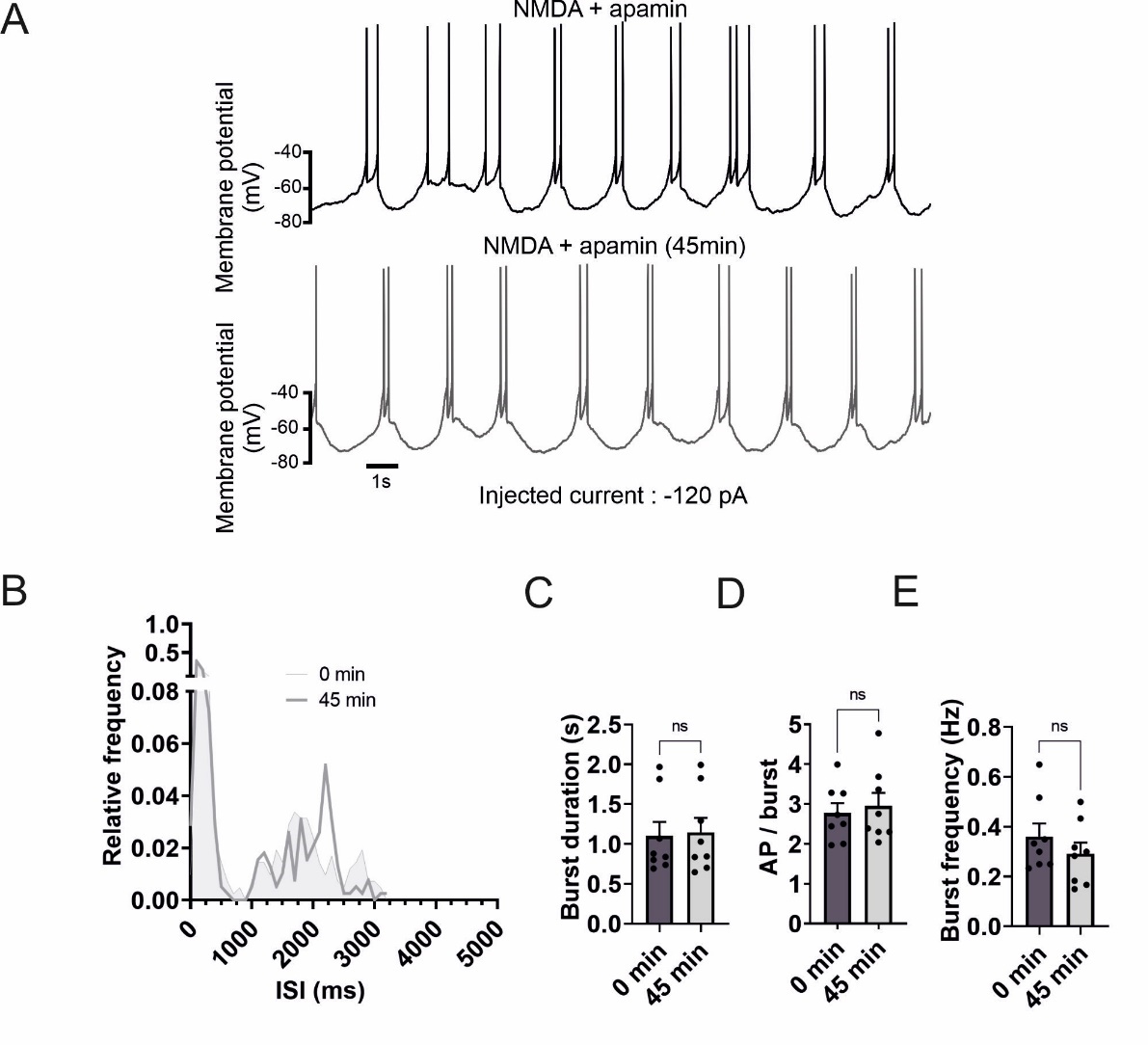
**Fig. S2. Stability of bursting activity through time**

**(A)** Voltage traces showing spontaneous action potential bursts in a SNc DA neuron in the presence of NMDA (30 μM) and apamin (300 nM, top trace) and after a recording time of 45 minutes (bottom trace). The dashed square at the right side of the voltage trace is a single burst with a higher magnification. **(B)** ISI histogram demonstrating the distribution of burst activity in control (light grey), and after 45 min (dark grey). **(C)** Summary plot showing burst duration in control and after 45 min. Control: 1.10 ± 0.18 s; after 45 min, 1.15 ± 0.18 s. Burst duration was not different in control and after 45 min (Wilcoxon signed rank test, ns). **(D)** Summary plot showing action potential per burst in in control and after 45 min. Control, 2.77 ± 0.25; after 45 min, 2.95 ± 0.33. No changes in the number of action potential per burst was observed (Paired T-test, ns) **(E)** Summary plot showing burst set rate in control and after 45 min. Control, 0.36 ± 0.05 Hz; after 45 min, 0.29 ± 0.05 Hz). Burst frequency was constant over time (Paired T-test, ns). Bars are means ± SEM for n=8.

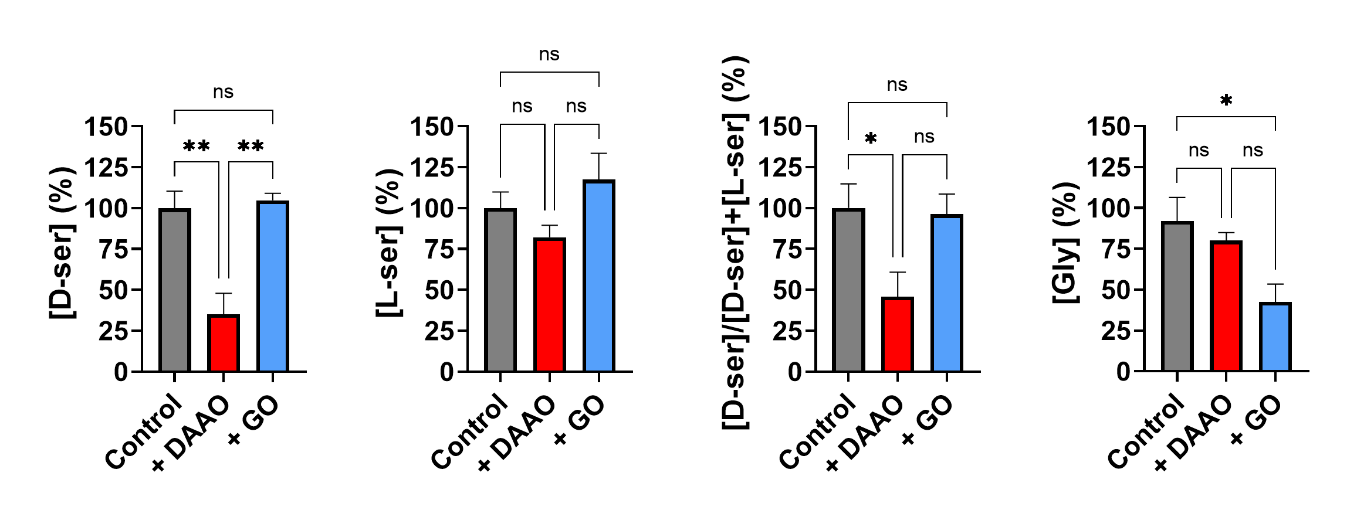

**Fig. S3. Specificity of DAAO and GO effects in the SNc.**

Summary plots showing the average concentrations of D-serine, L-serine (and the D-serine/total Ser ratio) and glycine determined by HPLC analysis of aCSF medium from SNc slices perfusion retrieved following experiments with control and samples treated with 0.2 U/mL of RgDAAO (+DAAO) or BsGO (+GO). Data are reported as percentage considering the control as 100% and as means ± SEM for n= 9 (control), n=6 (DAAO) and n=5 (GO); One-way anova (*P < 0.05; **P< 0.002). See text for details.

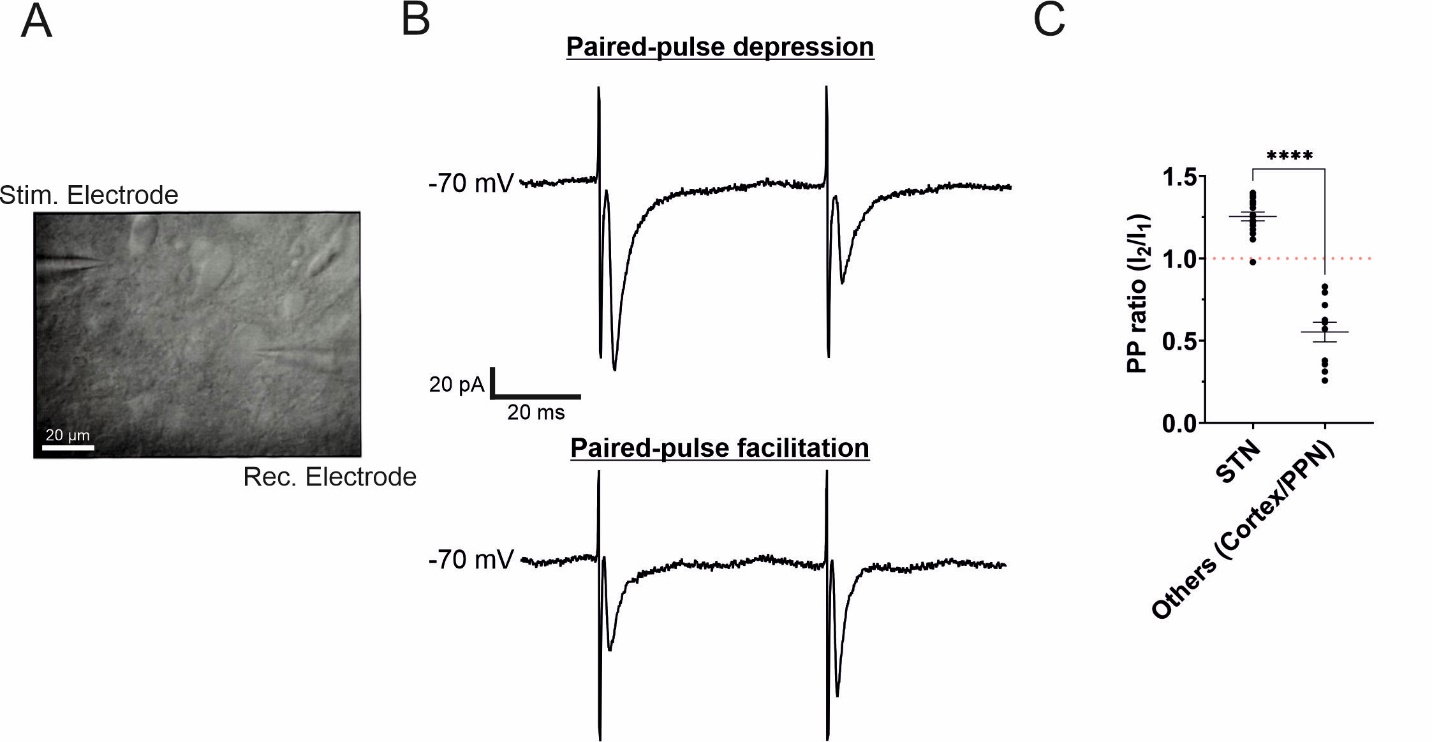

**Fig. S4. Paired-pulse ratio of afferent fibers originating from the STN**

**(A)** Infrared-Differential interference contrast photomicrograph of a DA neuron during whole-cell recording (right). The focal double-barrelled synaptic stimulating theta electrode was placed rostrally from the recorded neuron at approximatively 50-100 µm from the soma. **(B)** Current traces in response to a paired-pulse protocol (50 Hz) showing a paired-pulse facilitation when stimulating STN fibers (top) and paired-pulse depression when stimulating PPN fibers (bottom). Low stimulation levels were used (typically 20–80 µA) to evoke eEPSCs. **(C)** Summary plot showing the PPR as a function of the origin of the fibers stimulated. For STN fibers (n=17), the PPR ≥ 1 and for PPN fibers (n=11), the PPR ≤ 1 (STN, 1.26 ± 0.03 ; PPN, 0.55 ± 0.06) ; Unpaired T test, *****P* < 0.0001). This distribution is in accordance with the distribution obtained using optogenetic techniques (Beaudoin et al., 2018).

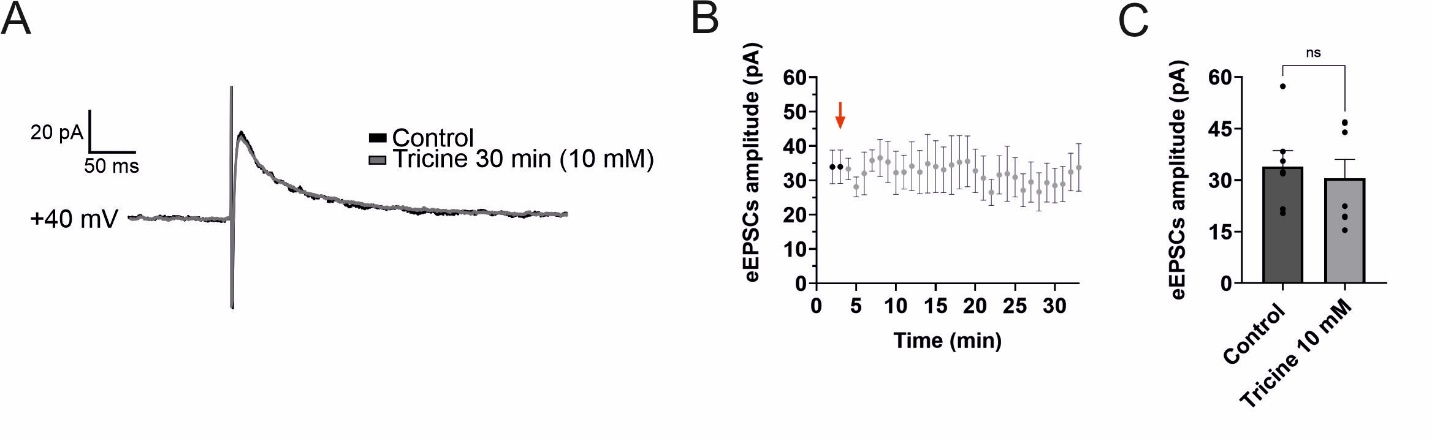

**Fig. S5. Effects of tricine for longer applications**

**(A)** Averaged eEPSCs (~30 events) recorded from SNc DA neurons at +40 mV in control conditions (black traces) and after incubation with Tricine (gray trace) for > 30 min. Low stimulation levels were used (typically 20–80 µA) to evoke eEPSCs. For each condition traces were recorded in the presence of picrotoxin (50 µM), CGP 55485 (1 µM), CNQX (10 µM), and strychnine (10 µM). **(B)** Averaged peak amplitude of NMDAR-eEPSCs immediately before (the two first points in black) and during the application of tricine for > 30 min. Note the constant amplitude of EPSCs during the application time. **(C)** Bar chart showing no difference (Wilcoxon signed rank test, ns) in the amplitude of the NMDAR-eEPSCs in control conditions (dark grey bar) and after 30 min in Tricine (grey bar). Control: 33.92 ± 4.66 pA ; Tricine, 30.57 ± 5.46 pA) ; Bars are means ± SEM for n=7.

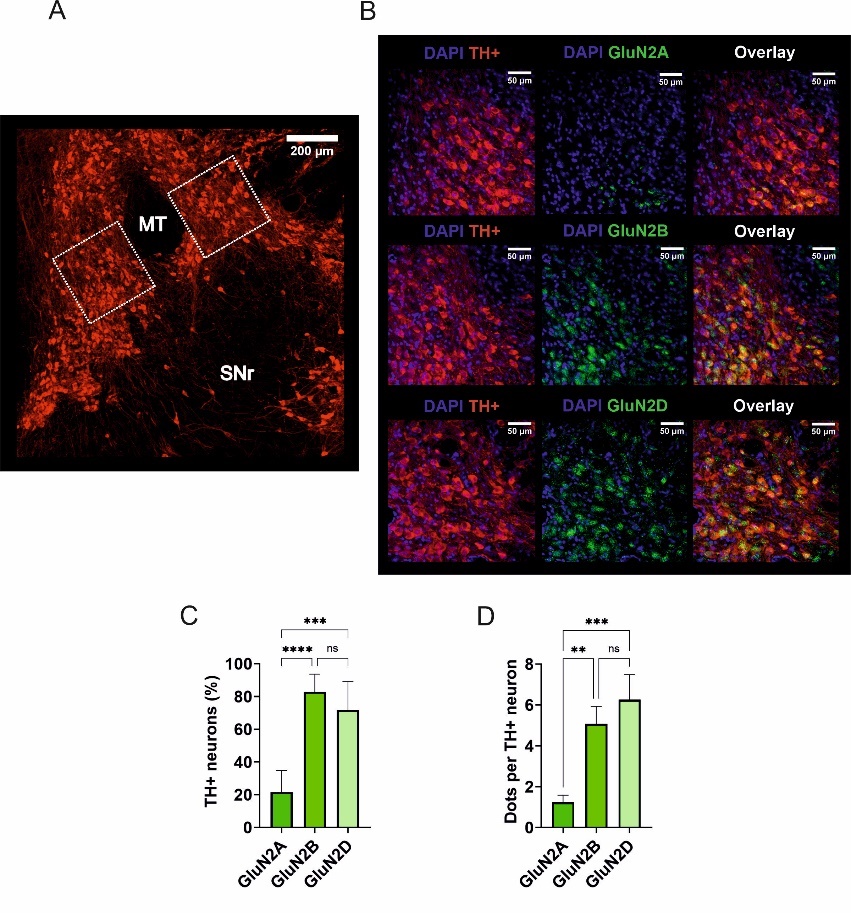
**Fig. S6. Expression of GluN2 NMDAR subunits in nigral DA neurons.**

**(A)** Light-micrograph of substantia nigra, showing immunolabeling for tyrosine hydroxylase (TH). MT refers to medial terminal nucleus of the accessory optic tract and has been used as an orientation point for the localization of the SNc. **(B)** Confocal light-micrograph of TH immunoreactivity (left), GluN2A immunoreactivity (top, center), GluN2B immunoreactivity (middle, center), GluN2D immunoreactivity (bottom, center), and overlay of GluN2A and TH (top, right), GluN2B and TH (middle, right) and GluN2D and TH (bottom, right); single confocal sections. **(C)** Summary bar graph showing the percentage of TH+ neurons expressing GluN2A, GluN2B or GluN2D. GluN2A: 21.54 ± 3.55 %; GluN2B, 82.76 ± 2.90 pA,; GluN2D, 71.71 ± 4.66 %. There are more GluN2B and GluN2D than GluN2A expressed in TH+ neurons (Kruskal-Wallis test followed by a post hoc Dunn’s test**,** *****P* < 0.0001, ****P* < 0.001 and ns) **(D)** Summary bar graph of the average number of fluorescent dots of either GluN2A, GluN2B or GluN2D in a TH+ neuron. GluN2A: 1.23 ± 0.0.35; GluN2B, 5.07 ± 0.85; GluN2D, 6.26 26 ± 0.1.22). There are more fluorescent dot of GluN2B and GluN2D than GluN2A expressed in TH+ neurons (Kruskal-Wallis test followed by a post hoc Dunn’s test**,** ***P* < 0.01, ****P* < 0.001 and ns) Bars are means ± SEM for n=7 (N=14).

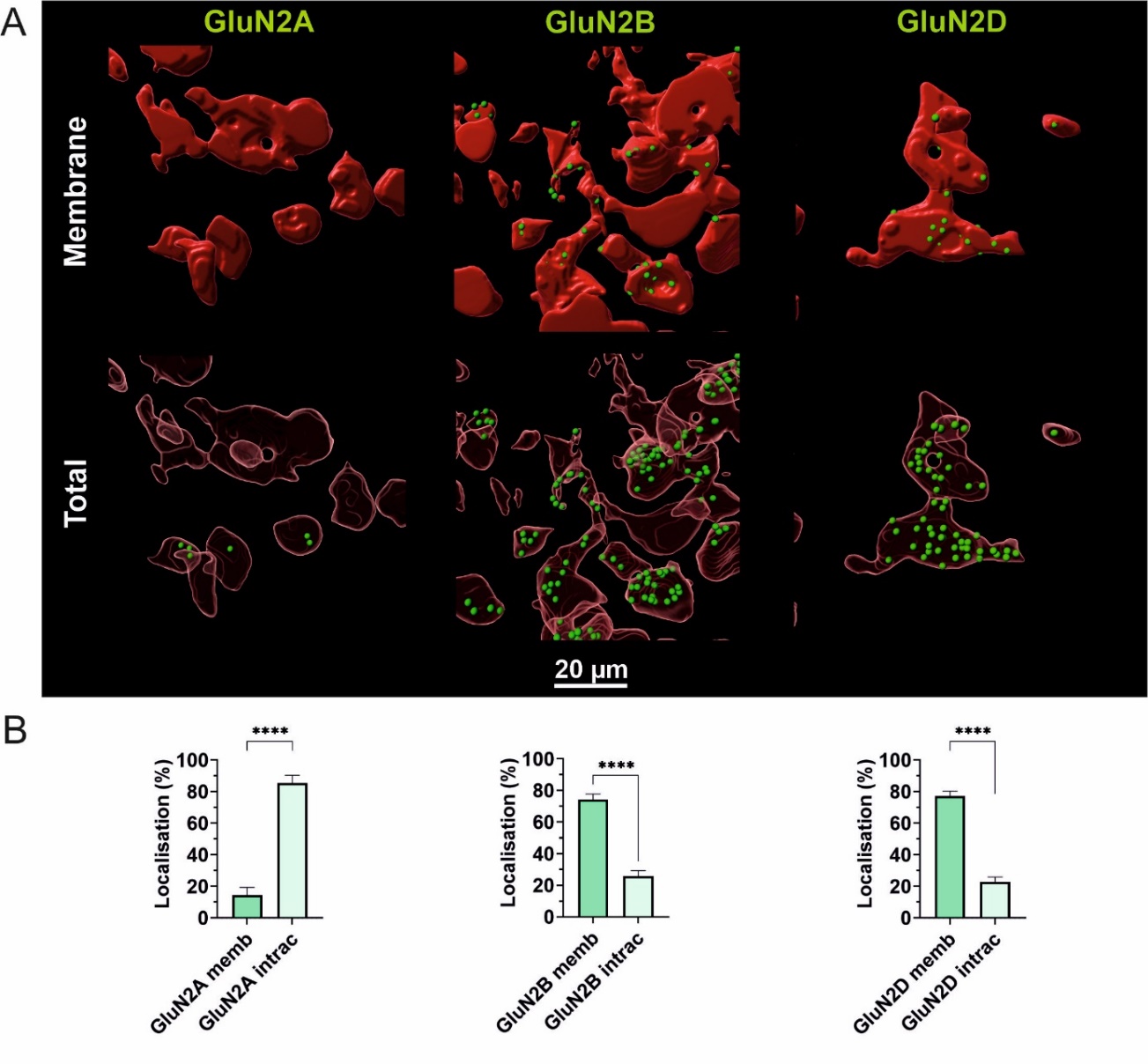
**Fig. S7. Spatial distribution of GluN2 NMDAR subunits in SNc DA neurons**

**(A)** 3D reconstruction of confocal images with DA neurons in red (TH+) and NMDAR subunits (green dots. Top panel represents the NMDAR subunits expressed only in the membrane of DA neurons and the bottom panel represents the expression of NMDAR subunits at the membrane, as well as, intracellularly. **(B)** Bar chart showing the localization of NMDAR subunits for GluN2A (left; in membrane, 14.52 ± 4.88 % ; intracellular, 85.48 ± 4.88 %, Mann-Whitney test, *****P* < 0.0001) , GluN2B (middle; in membrane, 74.24 ± 3.55 % ; intracellular, 25.76 ± 3.55 %, Mann-Whitney test, *****P* < 0.0001) and GluN2D (right; in membrane, 77.16 ± 3.14 % ; intracellular, 22.84 ± 3.14 %, Mann-Whitney test, *****P* < 0.0001). Bars are means ± SEM for n=7, N=35.

| **Experimental conditions** | **Burst duration (s)** | **AP/burst** | **Burst set rate**  **(Hz)** |
| --- | --- | --- | --- |
| Control  (N=6) | 0.90 ± 0.15 | 3.61 ± 0.48 | 0.36 ± 0.08 |
| DAAO  (n= 6) | 0.81 ± 0.12  ( p = 0.6171) | 4.34 ± 0.72  (p = 0.9519) | 0.26 ± 0.04  (p = 0.1376) |
| DAAO + D-Serine (100 µM)  (n= 6) | 1.41 ± 0.23  (p = 0.0124) | 6.16 ± 1.05  (p = 0.0014) | 0.23 ± 0.03  (p = 0.1684) |
| Control  (n= 8) | 0.92 ± 0.18 | 2.84 ± 0.22 | 0.47 ± 0.06 |
| GO  (n= 8) | 1.11 ± 0.23  (p = 0.7976) | 2.87 ± 0.29  (p > 0.9999) | 0.17 ± 0.05  (p = 0.0042) |
| GO + glycine (1 mM)  (n= 8)  **Numerical values are given as mean ± SEM** | 1.90 ± 0.51  (p = 0.0190) | 4.71 ± 1.10  (p = 0.0281) | 0.19 ± 0.05  (p = 0.0053) |

**Table S1. Effects of enzymatic treatment with DAAO and GO on bursting activity**

**Table S2. Effects of time and blockade of synaptic NMDARs on bursting activity**

| **Experimental conditions** | **Burst duration (s)** | **AP/burst** | **Burst set rate (Hz)** |
| --- | --- | --- | --- |
| Control 0 min  (N=8) | 1.10 ± 0.18 | 2.78 ± 0.25 | 0.36 ± 0.05 |
| Control 45 min  (n= 8) | 1.15 ± 0.18  (p = 0.8438) | 2.95 ± 0.33  (p = 0.3294) | 0.29 ± 0.05  (p = 0.1752) |
| synNMDAR blocked  (n= 6)  **Numerical values are given as mean ± SEM** | 1.43 ± 0.29  (p = 0.2824) | 2.20 ± 0.30  (p = 0.1636) | 0.35 ± 0.08  (p = 0.9106) |

**Movie S1. Surface expression of GluN2A subunits in nigral DA neurons.**

**Movie S2. Total expression of GluN2A subunits in nigral DA neurons.**

**Movie S3. Surface expression of GluN2B subunits in nigral DA neurons.**

**Movie S4. Total expression of GluN2B subunits in nigral DA neurons.**

**Movie S5. Surface expression of GluN2D subunits in nigral DA neurons.**

**Movie S6. Total expression of GluN2D subunits in nigral DA neurons.**

**.**
